# Megaplasmids associate with *Escherichia coli* and other *Enterobacteriaceae*

**DOI:** 10.1101/2025.09.30.679422

**Authors:** Allison K. Guitor, Shuai Wang, Owen T. Tuck, Brian Firek, Nadja Mostacci, April Jauhal, Lin-Xing Chen, Agata H. Dziegiel, Stephen Baker, Vu Thuy Duong, Alison E. Mather, Jukka Corander, Anu Kantele, Liat Shenhav, Markus Hilty, Michael J. Morowitz, Rohan Sachdeva, Jillian F. Banfield

## Abstract

Humans and animals are ubiquitously colonized by *Enterobacteriaceae*, a bacterial family that contains both commensals and clinically significant pathogens. Here, we report *Enterobacteriaceae* megaplasmids of up to 1.58 Mbp in length in infant and adult guts, and other microbiomes. Of 19 complete plasmid genomes, one was reconstructed from an *E. coli* isolate; others were linked to species of *Citrobacter* and *Enterobacter* via analysis of genome modification patterns. The detection of related plasmids in different *Enterobacteriaceae*, conjugation machinery, and more diverse modified motifs in certain plasmids compared to hosts suggests that these elements are self-transmissible, with a broad host range. The plasmids encode multi-drug efflux systems and potential secreted effectors. Up to 208 tRNAs are encoded and include sequence variants that may counter tRNA-centric defense mechanisms. Overall, the vast megaplasmid coding capacity may broaden host range, increase competitiveness, control invasion by other elements, and counter programmed cell death.

## Introduction

Plasmids are extrachromosomal elements that generally do not encapsulate and often can move between hosts. They are interesting from the perspective of lateral gene transfer, including medically relevant capacities such as antibiotic resistance and pathogenicity^1^. They may also encode host-relevant genes and hold high technological significance, for example, as genetic tools^2^. More broadly, they are integral and often overlooked components of microbial communities and evolution^3,4^.

The sizes of plasmid genomes can range from a few thousand base pairs to well over a million base pairs (i.e., megaplasmids - generally defined as plasmids >350 kb)^5^. Larger plasmids tend to have low copy numbers (PCNs), often a 1:1 ratio with the chromosome, and more metabolic capacities as compared to their smaller, higher copy-number counterparts^6–8^. Chromids and secondary chromosomes are large replicons with essential core genes. Secondary replicons in *Burkholderiales* with genomes of 2.1 - 3.36 Mbp were first suggested as megaplasmids, although an alternative classification as chromids or secondary chromosomes is now preferred^9–12^. Chromids, common in species of *Burkholderia*, *Brucella*, *Agrobacterium*, and *Vibrio*, tend to have plasmid-like replication and partitioning systems indicative of their origins from megaplasmids^5,13–16^. Secondary chromosomes are believed to have evolved from fragments of ancestral chromosomes^5,15^. Chromids and secondary chromosomes are often distinguished from megaplasmids by their essentiality and compositional signatures (e.g., GC content, codon usage) shared with the host’s primary chromosome^5,14,15^.

The best known megaplasmids are from *Rhizobium* (*Alphaproteobacteria*), with genomes up to 2.83 Mbp^17^, and 1.8 Mb plasmids of *Streptomyces clavuligerus* (*Actinomycetes*) with linear genomes^18^. In groundwater samples, the largest plasmids reported were circular and 2.96 and 1.74 Mb in length^19^. Surveys of plasmids from NCBI RefSeq and plasmid-specific databases representing diverse environments suggest a generally low prevalence of plasmids-like entities >1 Mb (0.03% - 1.17%) that mostly associate with *Alphaproteobacteria*^20–24^. Here, we report *Enterobacteriaceae* megaplasmids, including some that associate with *E. coli*. *E. coli* plasmids have been studied intensively, but the sizes of the best researched examples range from ∼5.8 kb (ColE1) to ∼100 kb (F (fertility) plasmid)^25,26^. In a collection of ∼4,500 circular *E. coli* plasmid genomes, the average size was 50 kb, although some clinically relevant IncF plasmids have ∼150 kb genomes^27^.

We show that *Enterobacteriaceae* megaplasmids are widely detected in human and animal microbiomes, and notably, that they can be present in the microbiomes of preterm infants, which are especially prone to colonization by *E. coli, Klebsiella, Enterobacter* and *Citrobacter*^28–30^. Previous detection of these sometimes highly abundant megaplasmids was likely prevented by assembly fragmentation and general focus on bacterial host genomes. Our analyses employed long-read and short-read sequencing along with manual curation to generate 20 confident, circularized genomes, of which 19 are fully validated and complete. *In silico* protein structure prediction was used to improve functional assignments, and methylation pattern analysis uncovered plasmid-host associations. Overall, we report new aspects of *Enterobacteriaceae* plasmid biology, with implications for the understanding of host bacteria that are especially relevant in human microbiomes.

## Results

### Large extrachromosomal elements in human infant gut metagenomic data

Long-read sequencing of a stool metagenomic sample from a preterm infant (INF1340011) identified an unusual large linear contig that was curated using Illumina reads into a circular genome of 1,583,849 bp in length (pMEGA1 - **Table 1**). As the genome does not encode rRNAs, ribosomal proteins, or core bacterial metabolic functions, we inferred that pMEGA1 is an extrachromosomal element (ECE).

**Table 1:** Listing of 4 complete plasmids (name, size, GC content, source). Completion was determined by mapping reads to a single contig and ensuring complete coverage as well as overlapping read coverage at the ends of the contig.

| Name | Size | GC content | Source | ORFs | tRNAs | Accession (sequencing technology) | Status |
| --- | --- | --- | --- | --- | --- | --- | --- |
| pMEGA1 | 1,583,849 | 26.11 | Preterm infant 1340011 8 weeks (USA) | 1633 | 119 | This study (Illumina + PacBio) | Complete & circular |
| pMEGA2 | 1,334,733 | 29.36 | Preterm infant 1330004 4 weeks (USA) | 1681 | 204 | This study (Illumina + PacBio) | Complete & circular |
| pMEGA3 | 1,373,843 | 29.57 | Preterm infant 1340011 4 weeks (USA) | 1691 | 170 | This study (PacBio) | Circular & confident |
| pMEGA4 | 1,415,043 | 25.98 | *E. coli* isolate from chicken rectal swab (Thailand) | 1483 | 108 | This study (Illumina + Nanopore) | Complete & circular |

Using long-read sequencing, we detected pMEGA1 in three other samples for infant 1340011 and found a different (MASH distance 0.24) large ECE from infant 1330004, for which manual curation generated a circular, complete genome of 1,334,733 bp (pMEGA2) (**Tables 1, S2**). Subsequently, we identified another PacBio sequence for a circular 1.37 Mb element (pMEGA3) in INF1340011, which did not have sufficient Illumina reads for curation to confirm all base calls (**Table 1**). The genome shares 98.3% ANI (88.6% aligned fraction) with pMEGA2 (**Table S2**). Interestingly, throughout the first 8 weeks of life, the gut microbiome of INF1340011 harbored pMEGA1 and pMEGA3 concurrently (**Figure S1)**. In both infants, these elements appeared as the gut microbiome shifted towards a community dominated by bacterial strains of the family *Enterobacteriaceae* (**Figures 1A, S1**). The maximum percentage of reads mapped to these ECEs in their respective infant gut samples was 15.74% for pMEGA1, 3.73% for pMEGA2, and 1.30% for pMEGA3 (**Figures 1A, S1; Tables S3, S4**).

**Figure 1:**
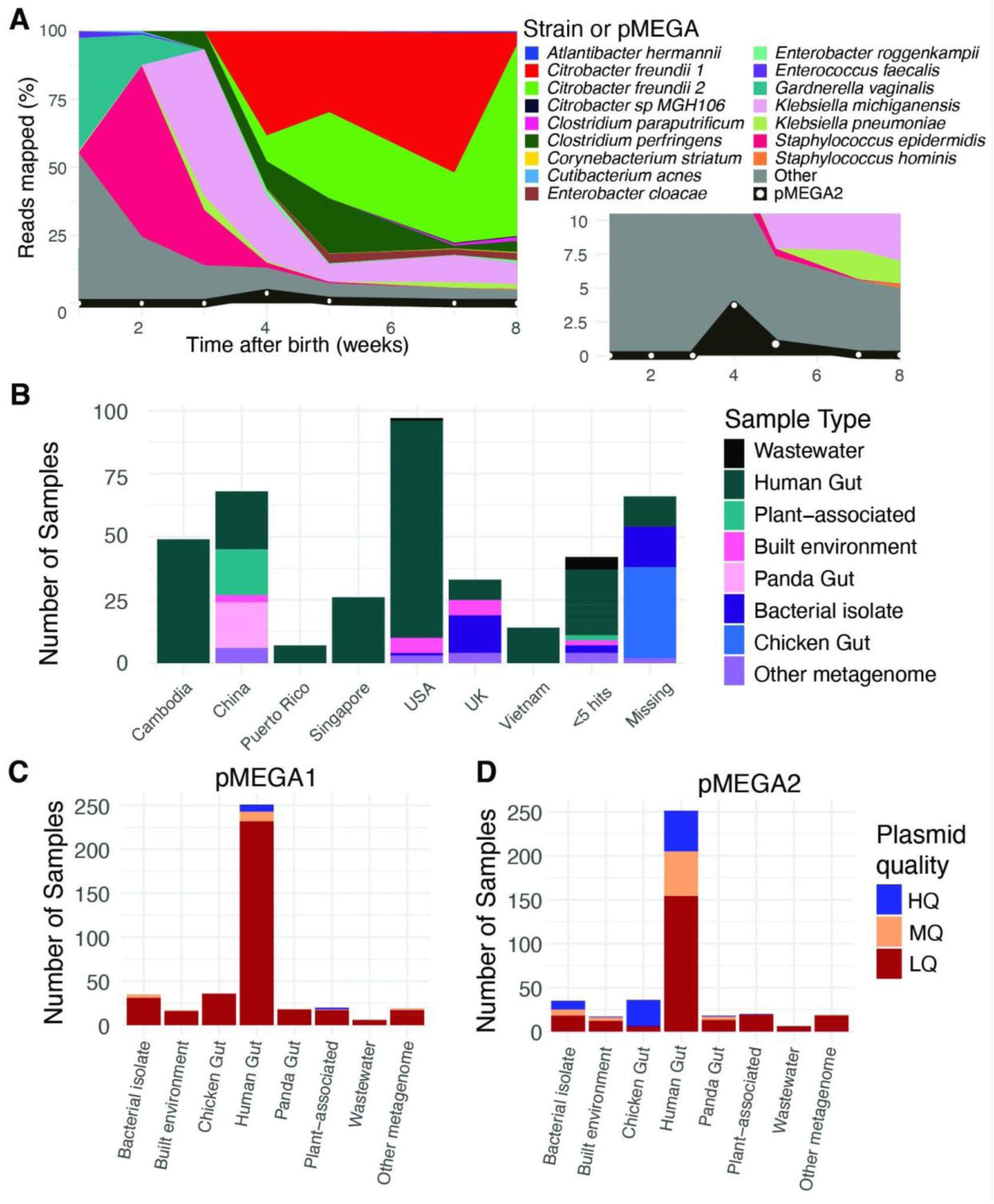
Identification of megaplasmids in the preterm infant gut microbiome and public data. **A)** The percentage of reads mapped to MAGs and pMEGA2 within INF1330004 was determined through Illumina read mapping. The “Other” category includes reads that did not map to MAGs nor pMEGA2. Inset shows the same plot but with y values from 0 to 10. **B)** Sources of samples with evidence of related megaplasmids, grouped by source location and sample type (Missing indicates no information in the SRA). Source types with fewer than 5 hits were grouped together. **C), D)** The number and quality of pMEGA1(**C**)- and pMEGA2(**D**)-like plasmids across different sample types. Assembled plasmids were classified as high-quality (HQ - blue) if they had ≥85% coverage of a pMEGA and ≤10 contigs and medium-quality (MQ - orange) with ≥85% coverage and 11-50 contigs. The remaining samples were classified as low-quality (LQ - red).

### Large extrachromosomal elements in bacterial isolates and various host-associated microbiomes

Through BlastP searches against NCBI, we identified an *Escherichia coli* isolate (strain G269_1R) from a rectal swab of a chicken from Bangkok, Thailand, with assembled fragments similar to pMEGA1^31^. After extensive manual curation of Oxford Nanopore long-read data with high-quality Illumina reads, we reconstructed a complete circular 1,415,043 bp genome (pMEGA4) that shares 95.8% ANI with pMEGA1 (85.0% aligned fraction) (**Tables 1, S2**). The host *E. coli* strain is a ST10 clone with a complete and polished 4,800,525 Mb genome and, in addition to pMEGA4, contains a 104 kb circular plasmid that we classified based on the presence of plasmid markers (i.e., *oriT*, *oriV*, relaxase, replication initiation protein, and partitioning proteins ParAB), and high similarity to other *E. coli* plasmids (NZ_CP070916.1, NZ_CP050045.1)^31^.

We surveyed > 400 metagenomic and bacterial isolate sequencing projects and found compelling evidence for almost 100 high-quality ECEs related to pMEGA1 and pMEGA2 from diverse sample types across multiple continents (**Figures 1BCD, S2; Table S5**). We detected elements in 10 infants for which we had time series data; in three infants, these elements persisted within the gut microbiome for months (**Tables S6**). Interestingly, these ECEs were also found in at least 23 *Escherichia coli* and 12 *Salmonella spp.* isolates (**Figure S2**). We predicted the PCN to range between 0.17 and 7.06, with *Salmonella spp*. showing a higher average PCN (2.79) than *E. coli* isolates (0.59) (**Table S5**). On average, the pMEGA1 and pMEGA2 sequences represent 0.69% and 2.98% of reads in samples, respectively.

We manually curated and confirmed the circularity of an additional 16 elements (**Table S7**). Given differences in size and GC content as well as ANI and MASH distance, we predict there are two distinct clades: Clade 1 represented by pMEGA1 and 4, and Clade 2 by pMEGA2 and 3 (**Figure S3AB; Table S2**). There is some synteny across the elements, but there is also evidence of extensive rearrangements (**Figure S3C**).

### Identification of pMEGA elements as megaplasmids

Given that ECEs are dominated by relatively novel proteins, we used *in silico* protein structure prediction in addition to other annotation tools to predict functional annotations for pMEGAs 1-4 (**Methods**). We did not identify proteins that would suggest classification as bacteriophages. Only a few genes encode proteins directly related to core bacterial metabolism, suggesting that these elements are not secondary chromosomes/chromids (**Table S8**). Further support against classification as a chromid or secondary chromosome is the difference in GC content between pMEGA4 (25.98%) and its *E. coli* host genome (50.83%).

Consistent with the identification as megaplasmids, we identified genes for the partitioning proteins (ParC,E (pMEGA4_1,2) components of Topoisomerase IV) and ParF (pMEGA4_1239), a ParA family NTPase of the Type-Ib partitioning system typically co-located with ParG in low copy number plasmids^32^. Although a ParG homolog was not identified, we identified PadC, a protein with a ParB-like nuclease domain (pMEGA4_1030) known to interact with ParA.

We also identified most components of a Type IV secretion system within the first 80 kb of the genome (e.g., (Vir/Dot/Icm proteins), two relaxases (pMEGA4_58, 1394), and two subfamilies of replication initiation (Rep) proteins (**Figure S4AB; Tables S8, S9**). The putative relaxases are most similar to the MOB_F_ family, which includes many large and conjugative plasmids, and clade with plasmids from diverse bacterial hosts (**Figure S4B**)^33^. An intergenic AT-rich region adjacent to a relaxase (pMEGA4_1394) could serve as an origin of transfer (OriT), but this was not identified as such by oriTFinder2^34^. The predicted Rep proteins (pMEGA4_629, 1423) share <31% amino acid identity with bacterial sequences and the predicted structures are similar to those of RepC from an IncQ plasmid in *Salmonella enteritidis* and TrfA from an IncP plasmid in *Escherichia coli* (**Figure S4C**). However, the Rep proteins form distinct phylogenetic clades, and thus these plasmids are only distantly related to IncQ and IncP plasmids (**Figure S4A**). One origin of replication (OriV) with five 18 bp direct repeats was detected upstream of pMEGA4_1423, and another possible iteron-containing locus with less conserved repeats occurs downstream of pMEGA4_629 (**Figure S4D**)^35^. These features, and a type IV coupling protein (T4CP pMEGA4_20), indicate that these are self-transmissible conjugative megaplasmids^33,36^.

### *Enterobacteriaceae* host these megaplasmids

pMEGA4 was reconstructed from an *E. coli* isolate, and pMEGA17 and near-identical pMEGA18 were reconstructed from two impure *E. coli* isolates (**Table S7**). Incomplete megaplasmid genomes were reconstructed from isolates of *Salmonella enterica* and *E. coli*. We attempted to identify the pMEGA1, 2, and 3 hosts using DNA crosslinking applied to stool samples from two infants, but the results were inconclusive. While most proteins (>66%) in pMEGA1-3 have an unknown taxonomic classification, ∼8% have sequence similarity to members of the order *Enterobacterales* (**Table S7**).

To further identify the pMEGA1-3 hosts, we used abundance-based correlation analysis. The strongest correlation in abundance for pMEGA1 was with *Enterobacter cloacae* 1 (0.92), for pMEGA2, *Citrobacter freundii* 1 (0.88), and for pMEGA3, *Klebsiella michiganensis* 1 (0.9) (**Table S10**). We also investigated DNA modification patterns using PacBio HiFi reads and compared potentially modified motifs shared between pMEGAs, coexisting bacterial genomes, phage, and plasmids (**Figures 2, S5**). Strong overlap in modified motifs between pMEGA2 and the *Citrobacter freundii* 1 genome supports the correlation-based host prediction. For pMEGA1, which only shares motif patterns with pMEGA3, the results are inconclusive (**Figure S5**). pMEGA3 has overlapping motifs with a *Citrobacter freundii* and an *Enterobacter cloacae* strain. The pMEGAs have a greater number and diversity of modified motifs compared to bacterial genomes within the same microbiome sample. Paralleling this, pMEGA4 encodes 21 methyltransferases (MTs), whereas the host *E. coli* genome encodes only 6 (**Table S11**).

**Figure 2:**
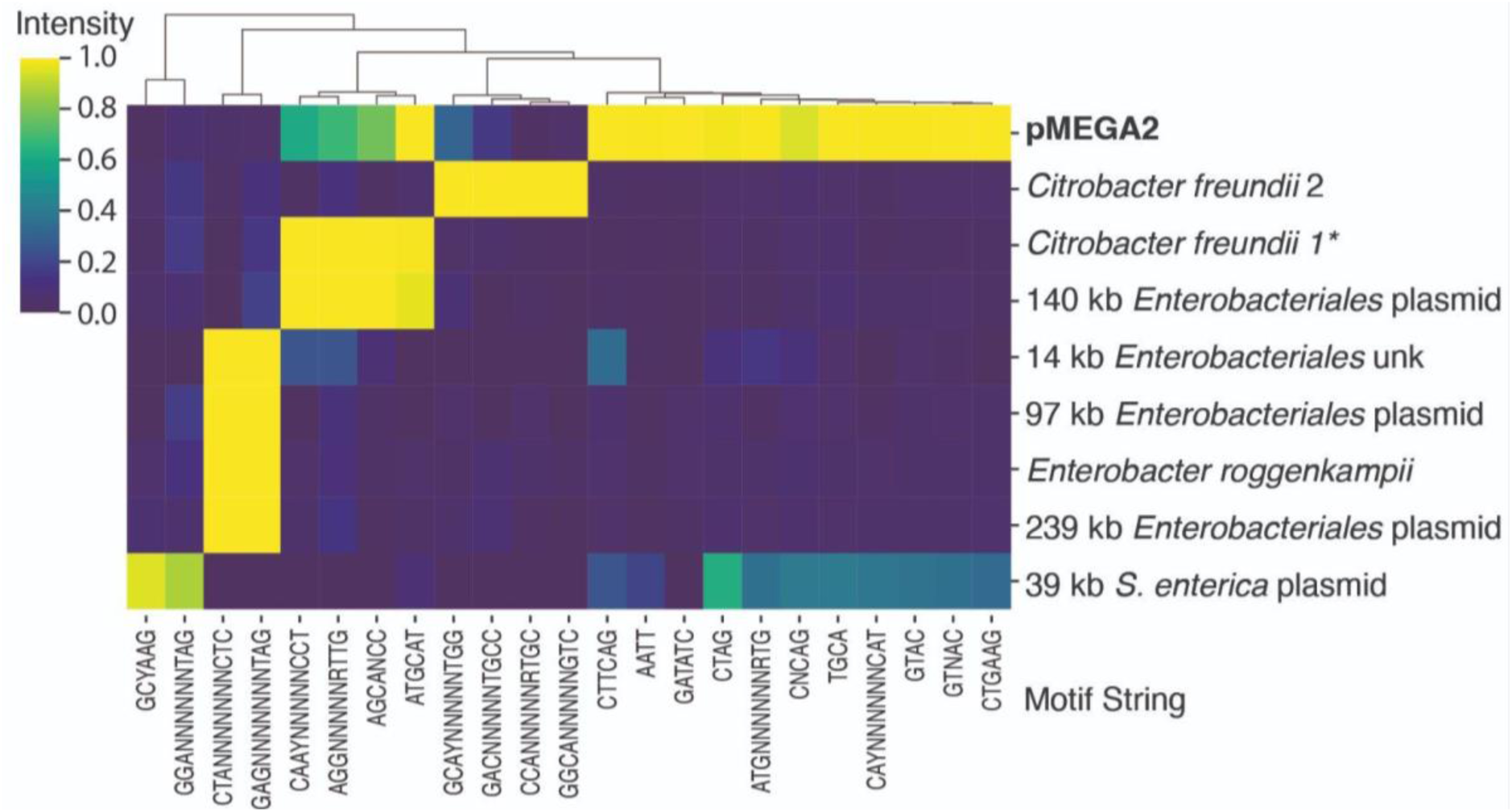
A *Citrobacter freundii* is the host of pMEGA2. Modified bases were predicted in the PacBio HiFi reads of infant 1330004 at week 4. The columns represent the motifs predicted to contain modified bases, and the rows represent genomes. *Likely host of pMEGA2 based on overlap in modification patterns.

pMEGA4 encodes a type-I-Fv CRISPR-Cas system with a complete Cas1-Cas2 integrase, a Cas3 helicase-nuclease effector, and a compact Cascade surveillance complex likely encoding only five subunits (**Figure S6A-D**)^37^. Components of these systems are not completely conserved in all pMEGAs. Consistent with active adaptive immune activity, all pMEGAs contain CRISPR arrays with between 27 and 220 spacers (**Table S7**). No CRISPR spacers from the pMEGAs targeted coexisting bacterial genomes, and vice versa. However, pMEGAs share some CRISPR spacer sequences with other pMEGAs from the same clade (but not between clades), and a subset of plasmid CRISPR spacers target Cas genes of other pMEGAs (**Figure S6E**). Many spacers of pMEGAs target plasmids from *Enterobacteriaceae*, consistent with the role of plasmid CRISPR loci in plasmid-plasmid competition (**Figure S6E**).

We used CRISPR spacers from databases and metagenomes to search for targeting of pMEGAs that may be indicative of megaplasmid host associations^38,39^. We identified 12 non-redundant spacers (100% match) from 33 metagenomes and an isolate of *Proteus cibarius* (*Enterobacteriaceae*) that target regions in at least one pMEGA (expectation for random match of ∼e^−11^) (**Table S12**). The *Proteus* isolate spacer targets a tRNA_Met_ locus that is conserved in pMEGA1, 4, and 20^40^. Five of the 11 metagenome-derived spacers also target tRNAs. Overall, based on isolate source, protein similarity, co-occurrence patterns, CRISPR targeting and methylation analysis, we conclude that various *Enterobacteriaceae* host the megaplasmids.

To put these megabase-scale plasmids into context, we analyzed the PLSDB^21^ and IMG/PR^22^ databases and estimated that of the >770,000 plasmids, there are 667 non-redundant >1 Mb genomes, all but four of which are from isolates. Only 30 were detected in *Gammaproteobacteria* **(Figure S7).** In addition, nine circular and five linear >1 Mb secondary replicons of *Enterobacteriaceae* genomes are listed in NCBI^23,41^. Given similar GC content compared to the primary chromosome and genes for many core functions (including multiple rRNAs in some cases), we suspected that they are not megaplasmids (**Supplementary information; Table S13**). We assayed hypothetical gene content of the public putative *Enterobacteriaceae* megaplasmids and found essentially no differences compared to the main chromosome (24.9% vs 23.2%) (**Table S13**). This is in stark contrast to the pMEGAs which feature ∼ 80% hypothetical genes, compared to 21.1% for the *E. coli* host of pMEGA4 (**Table S13**). We conclude that these *Enterobacteriaceae* sequences are secondary chromosomes or parts of the main chromosomes.

### Clades 1 and 2 differ in their multi-copy protein sets

Of 2,162 protein subfamilies (SF), 438 occur in all of the 20 pMEGAs (**Table S9; Figure 3A**). These conserved proteins are involved in transcription, translation, replication, protein folding, defense, pilus formation, and secretion systems. Many are annotated as proteases, phosphatases, transport-related, methyltransferases, chemotaxis proteins and proteins involved in NAD metabolism and acetylation/deacetylation (**Table S9; Figure 3A**). 336 of the 438 ubiquitous proteins occur only once per genome, typically in similar genomic locations within clades (but organization differs between clades; **Figure 3C**).

**Figure 3:**
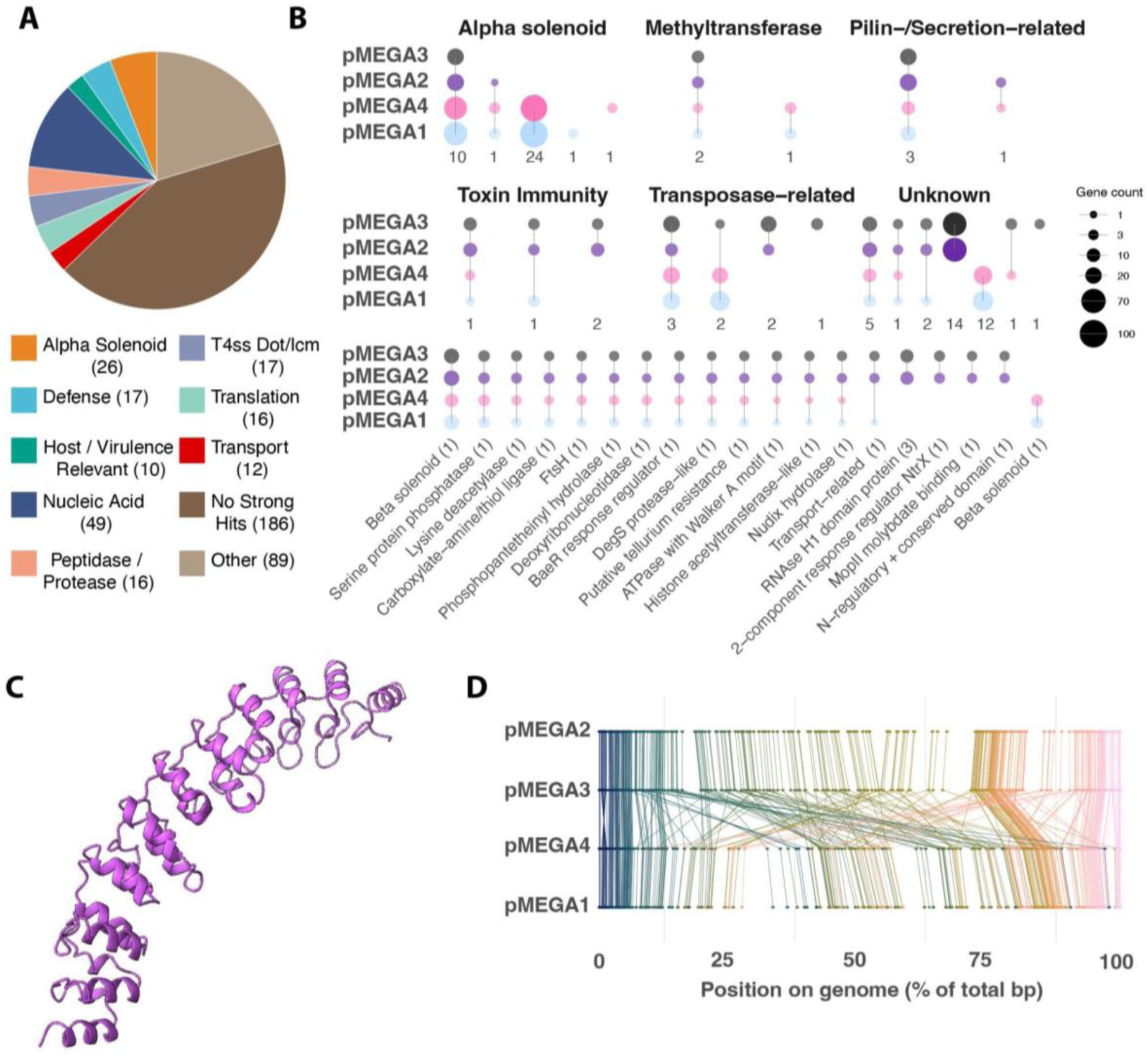
Conserved protein functions across two clades of megaplasmids. **A)** Distribution of functional annotations of the 438 conserved protein subfamilies across pMEGAs1-20. Annotations with less than 10 instances were grouped into the “Other” category. Numbers represent the number of protein families in each category. **B)** Multicopy protein subfamilies across pMEGA1-4. Circle size corresponds to the number of genes within each category. The number below each column or at the end of each annotation in the Other row, indicates the number of subfamilies in that category. **C)** Examples of alpha solenoid-containing protein pMEGA4_362 (median pLDDT 95.06) whose domain is annotated as an Ankyrin repeat. **D)** Alignment of pMEGA1-4 showing the locations of the 336 single copy protein subfamilies conserved across pMEGA1-20 as a percentage of the total length of the genome. Each protein subfamily is coloured based on their gene order in pMEGA2.

Predicated on the idea that multicopy proteins (here defined as 3 or more copies/genome) may be important to the functioning of megaplasmids, we counted the instances of each subfamily in the pMEGA1-4 genomes (**Table S14**; **Figure 3B**). Of the 10 subfamilies that are multicopy in all four megaplasmids, the most prevalent are beta-solenoid structures (SF0950: 10 - 16 per genome), possible bacterial shufflons/pilin proteins (SF2096: 7 - 16 per genome), TnpB proteins (SF1483: up to 11 per genome), and protein phosphatases (SF1635). pMEGA1 and 4, and to a lesser extent, pMEGA2 and 3, have numerous proteins from many subfamilies with predicted alpha-solenoid structures. The alpha-solenoids are of varying lengths and are encoded by >300 genes (∼20% of the proteome) of pMEGA4 (**Figure 3BC**). Many had very low-scoring hits to structures in the PDB, and those with higher-scoring matches were mostly to Designed Ankyrin Repeat Proteins (DARPins) that are of interest as protein-based therapeutics^42^. Multicopy proteins that are only highly prevalent in pMEGA1 and 4 are serine recombinases (SF2001: TnpA - up to 23 copies) and their accompanying endonucleases, and DNA methyltransferases (SF0095), whereas novel proteins, possible immunity-related proteins, and three distinct RNAse H1 subfamilies are only highly prevalent in pMEGA2 and 3 (**Figure 3B**).

Some multi-copy protein subfamilies are clustered in the genomes (**Table S14)**. Two to four copies of proteins in the same subfamily occur sequentially in 58 and 33 instances in pMEGA1 and 4, respectively. In pMEGA2 and 3, two to eleven genes of the same subfamily occur sequentially 92 and 110 times, and there are eight RNAse H1 subfamily proteins encoded almost consecutively. In addition to the conserved RNAse H domain, these proteins have diverse additional regions/domains, sometimes positively charged (possibly involved in DNA binding), but for which functions could not be assigned.

Copies of SF0552, proteins similar to a toxin immunity protein, are clustered in pMEGA1, 2 and 3 (**Figure S8A**). Encoded immediately upstream is a protein with an AHH domain (SF1803), commonly found in secreted nuclease toxin effectors (**Figure S8BDE**)^43^. When folded together, the putative immunity protein is confidently predicted (chain_pair_iptm values 0.91-0.92) to interact with the AHH domain (**Figure S8C**). The cluster is reminiscent of poly-immunity loci associated with polymorphic toxins^44–46^. pMEGA2 and 3 have an additional region with the putative toxin SF1803 encoded upstream of a cluster of possibly immunity proteins SF0823. To screen for other toxin/poly-immunity loci, we analyzed proteins directly upstream of multicopy clustered proteins to identify toxin domains. We found SF0417 as a possible ADP-ribosyltransferase upstream of 3 and 5 copies of SF0062 in pMEGA2 and 3, and a possible protease (SF0355) upstream of 3 copies of SF0744 in pMEGA2. Poly-immunity appears more common in pMEGA2 and 3 as compared to pMEGA1 and 4.

### Functional overlap and differences between pMEGA4 and *E. coli* host

Given that pMEGA4 originated from an *E. coli* isolate, we compared the predicted genetic repertoires and proteomes of this megaplasmid to its host in detail (**Table S15**). Of note, only 9 proteins shared >70% amino acid identity to host proteins (**Table S16**). pMEGA4 encodes numerous proteins involved in nucleic acid manipulation, including polymerases, helicases, nucleases, proteins involved in recombination and nucleotide interconversion (e.g., GMP->GDP) and over 20 translation-related proteins (**Table S8**). These functions are also encoded on the *E. coli* genome.

Possibly involved in homeostasis and quality control, the pMEGA4 genome encodes the protease ClpP (pMEGA4_187) and associated proteins^47–50^(**Figure S9AB**). The genome encodes three copies of proteins with high structural similarity to FtsH (pMEGA4_57, 182, 1364) and three proteins similar to HflK/C that multimerize into functional microdomains (**Figure S9CD**)^51,52^. The *E. coli* genome encodes ClpP and associated proteins, HflK/C, as well as one copy of FtsH.

In addition to a role in conjugation, T4SSs may be involved in effector protein delivery and virulence (similar regions occur in pMEGAs1,2 and 3)^53^. The *E. coli* host genome encodes components of Type II and Type VI secretion systems, including VgrG and Hcp, neither of which are identified in pMEGA4^54^. Most interesting, and seemingly not found in the *E. coli* host, is a protein with structural homology to InvG^55^ (**Figure 4ABC**).

**Figure 4:**
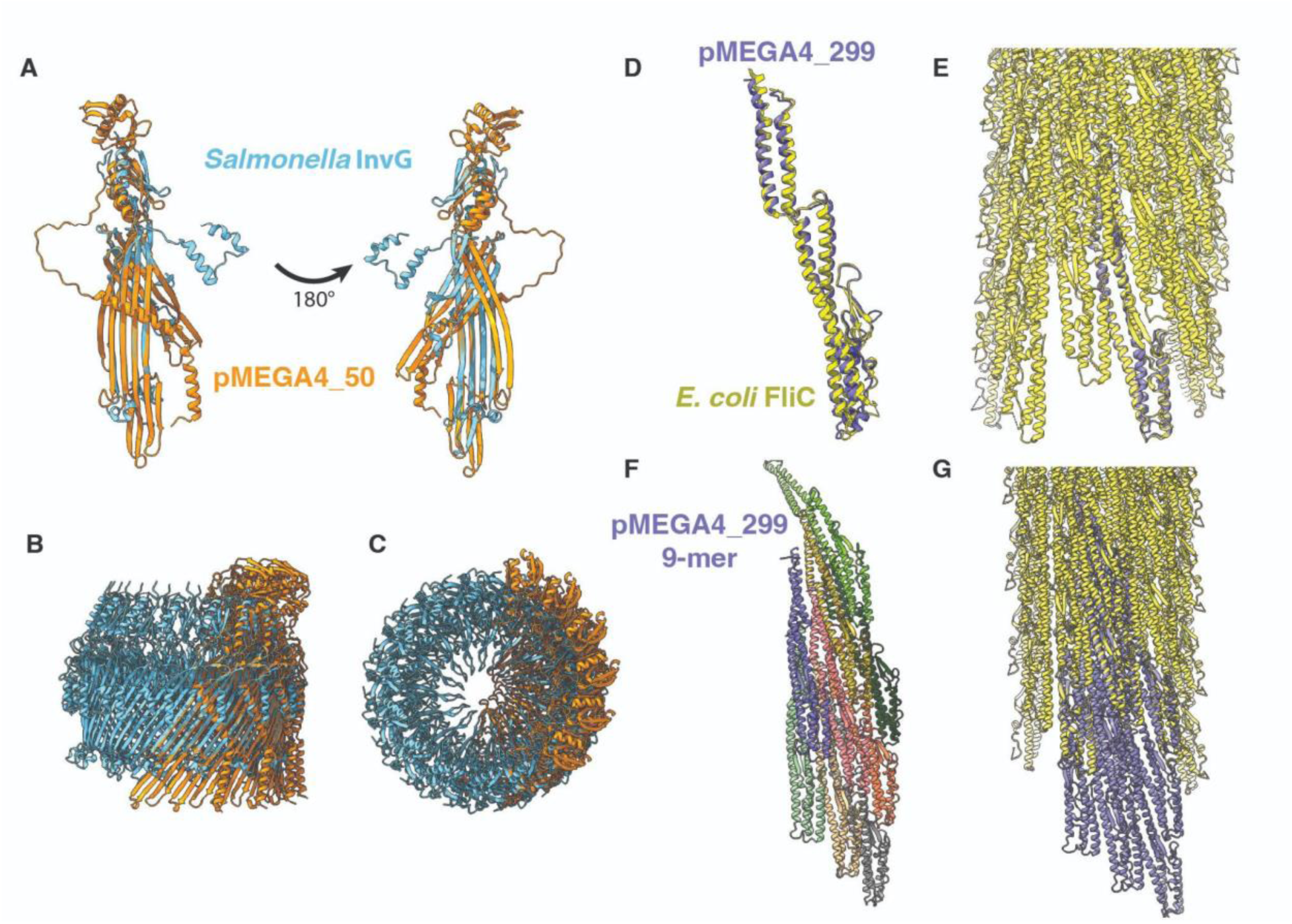
pMEGA4 encodes mechanisms related to secretion and virulence. **A)** Predicted structure of a monomer of pMEGA4_50 (median pLDDT 82.0) aligned to *Salmonella* pathogenicity island 1 injectisome protein InvG (PDB: 6PEE). Predicted hexamer of pMEGA4_50 aligned to the 15-mer complex of InvG shown from the side (**B**) and top (**C**). **D)** Predicted structure of monomer of pMEGA4_299 (median pLDDT 93.91) aligned to *E. coli* K12 flagellin monomer (PDB: 8XCM) and **E)** full filament core. **F)** Predicted homo-nonamer structure of pMEGA4_299. **G)** Alignment of predicted 9-mer of pMEGA4_299 with flagellar core of *E. coli* K12 (PDB:8XCM)^175^.

Also encoded in the pMEGA4 genome are at least four proteins (pMEGA4_1281, 90, 966, 977) with a von Willebrand factor A–like domain (vWA), which is commonly associated with adhesins. The *E. coli* genome encodes two proteins with vWA domains. Another protein seemingly redundant with the host *E. coli* repertoire, pMEGA4_299 encoded *fliC,* multimerized nicely into a typical flagellin structure. This appendage is involved in adhesion, motility, virulence, and immune stimulation^56,57^ (**Figure 4DEFG**).

pMEGA4 encodes 105 verified tRNAs, including at least one for each canonical amino acid with 91 unique tRNA sequence types (87%), whereas the *E. coli* genome encodes 86 of which 49 are unique (57%) (**Table S17**). The tRNAs are mostly arrayed in blocks, generally with ∼6 to ∼ 25 nucleotide (nt) gaps between them (**Figure 5A**). Only two blocks (with 7 and 4 tRNAs) occur in the *E. coli* genome. Within the set of complete pMEGA genomes, there are up to 208 tRNAs, with statistically significant enrichment in Clade 2 (average 199 ± 9) compared to Clade 1 (average 113 ± 8).

**Figure 5:**
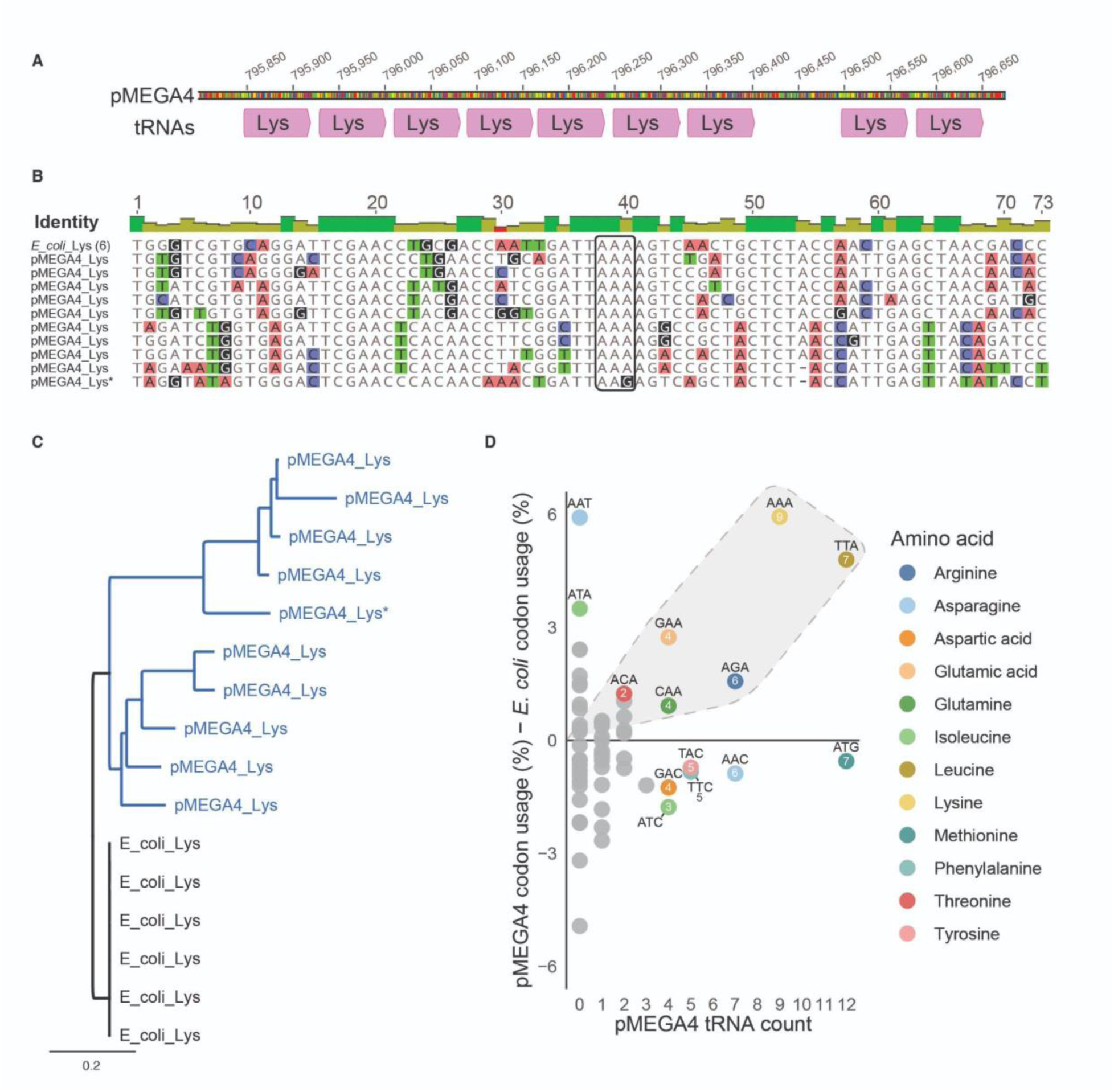
tRNAs of pMEGA4 and *E. coli* host. **A)** Nine of the ten pMEGA4 tRNAs for LYS are encoded in a single block. **B**) Nine of the ten pMEGA4 LYS tRNA genes have the same (AAA) codon, however the sequences differ. **C)** Phylogenetic tree constructed using LYS tRNAs for pMEGA4 and host *E. coli*, illustrating pMEGA4 sequence diversity yet conservation of the host sequences. The asterisk indicates the pMEGA4 AAG codon. **D**) Some high tRNA counts are predicted by more extensive use of that codon in pMEGA4 compared to *E. coli* (dots in dashed outlined region). However, other highly represented codons do not follow this pattern. Numbers associated with each point (colored by amino acid) indicate the count of unique tRNA sequences.

For the tRNAs for the most highly represented amino acids (Leu, Lys, (Met)), almost all have the same codon. For Lys, 9/10 are encoded in one block and all have the same codon (AAA) despite being quite divergent in the rest of their sequences (**Figure 5B**). In contrast, the six *E. coli* tRNA Lys are all identical (**Figure 5C**). For 11 other amino acids, all tRNAs for that amino acid are in a single block. In contrast, in the *E. coli* host genome, there is only one tRNA (Gln) where all instances occur within a block (two codon types). In some cases, different tRNA types alternate pseudo-regularly (sometimes with three or more different tRNAs per cluster).

A commonly proposed explanation for mobile element tRNAs is compensation for differences in their codon usage compared to that of their host. We find that some codons much preferred by pMEGA4 compared to the host *E. coli* have more highly represented tRNAs (e.g., AAA, TTA), but this is not the rule (**Figure 5D; Table S18**). Some less commonly used tRNA codons (e.g., ATG, AAC) are highly represented, and some much more highly used codons (e.g., AAT, ATA) are not represented in pMEGA4. Notably, there is high sequence diversity associated with all highly used codons (**Figure 5D**).

### pMEGA4 may confer stress protection, aid in host competition and antibiotic resistance

pMEGA4 encodes a protein that forms a ferritin-like cage for DNA protection during starvation (pMEGA4_661; also identified in *E. coli*) and two small alarmone hydrolases that regulate the stringent response in a similar way to *gppA* found in the *E. coli*^58,59^. Further supporting defense against external challenges are genes for redox homeostasis, nucleic acid repair, oxidative stress, heat shock, and genes that form, maintain and repair disulfide bonds. Again, these functions appear redundant with those encoded in the *E. coli* genome. pMEGA4 encodes the outer membrane protein SlyB (also encoded in the host genome), which is a lipoprotein that forms a 13-mer pore (**Figure S6C**). The PhoPQ regulon is responsible for the expression of SlyB in response to external stress^60^. We also identified three near-consecutive genes for tellurium resistance (TerD, not identified in *E. coli*).

Possibly in response to bacterial competition, pMEGA4 encodes a protein implicated in immunity to effectors (pMEGA4_953)^61^, an amidase antitoxin (pMEGA4_699), and a hydrolase against ADP-ribosylating toxins (pMEGA4_1469^62^). These antitoxins appear without their toxin counterparts. Conversely, there are at least two proteins annotated as the lipoprotein toxin entericidin B that are separate on the plasmid and do not appear co-localized with an antitoxin (SF1615)^63^. The *E. coli* host genome encodes multiple Type II toxin/antitoxin systems (ie. MazEF, HipAB) and many possible secreted effectors/toxins ^64–66^.

In addition to the CRISPR-Cas system described earlier, pMEGA4 encodes putative defense system proteins structurally similar to the Upf1 family of RNA helicases (pMEGA4_338,520; detected by PADLOC as Mokosh TypeII systems), and a Ser/Thr kinase annotated as a PD-T4-6 system (pMEGA4_91)^67–69^. The *E. coli* genome encodes 7 single-gene phage defense candidate systems, 2 Hma (Helicase, Methylase, ATPase) defense systems, HEC, hhe, and a CRISPR array^70^.

The *E. coli* host of pMEGA4 encodes the extended-spectrum beta-lactamase (ESBL) *bla*_CTX-M-55_ on the abovementioned 104 kb plasmid, along with genes for quinolone (*qnrS1*), chloramphenicol (*catII*), and tetracycline resistance (*tetR* and *tetA*)^31^ (**Table S19**). Few antibiotic resistance mechanisms were detected in pMEGA4 (**Table S19**). However, this function may be conferred by 10 genes for transporters, including MacAB- and AcrAB-TolC efflux systems that were identified based on sequences and predicted structures^71^. Possibly relating to antibiotic resistance, pMEGA4 contains an outer membrane protein/porin implicated in carbapenem import (pMEGA4_670), and a methyltransferase that may activate a peptide antibiotic (pMEGA4_515).

### Panda gut metatranscriptomic data reveals broad gene expression

We identified a public metatranscriptomic dataset from a panda gut microbiome with both a clade 1 and a clade 2 megaplasmid (SRR10902853). As there was no DNA data, we mapped the reads to pMEGAs 1 - 4 competitively and detected extensive read recruitment to all four genomes. By far the highest recruitment was to pMEGA4 with almost one million reads, but over 128,000 reads were recruited to pMEGA3. From the transcripts, we assembled 1,613 contigs > 5 kb in length, with a maximum length of 145 kb (4 of the 5 longest scaffolds were from *E. coli* and *Klebsiella* phages), and a 109 kb scaffold with similarity to pMEGA4. 75 contigs > 5 kb mapped uniquely to either clade 1 (pMEGA4) or clade 2 (pMEGA4) genomes with no overlap of contigs between clades. For the clade 1-like plasmid, we recovered 940,590 bp with ANI of 99.52% across 63.81% of pMEGA4. We used the transcripts to curate the 109 kb sequence from the clade 1 genome, primarily closing two scaffolding gaps (**Figure 6A**). For the clade 2-like plasmid, we recovered 563,270 bp with ANI of 98.70% across 37.93% of pMEGA3.

**Figure 6:**
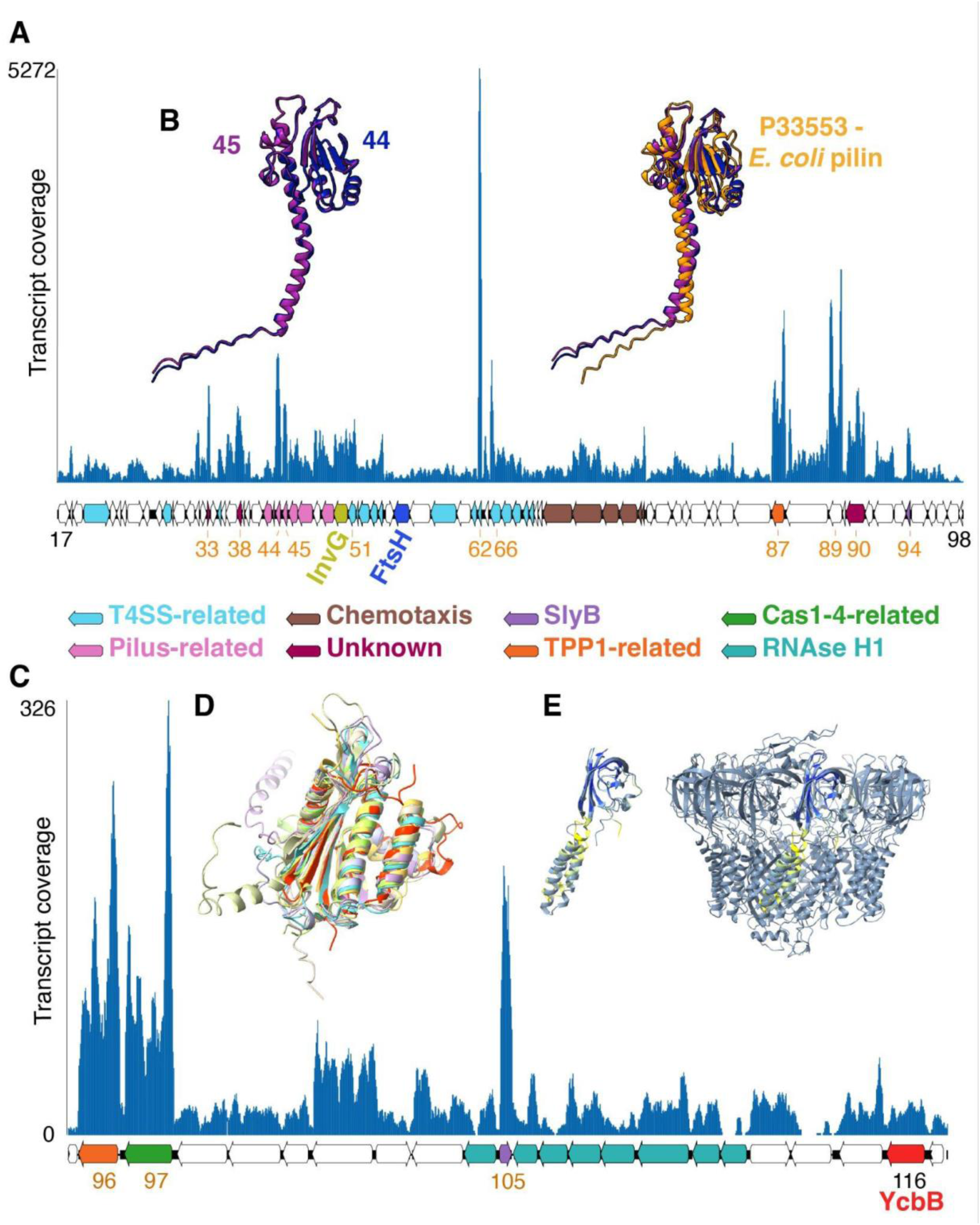
Public data (SRR10902853) from a panda gut metatranscriptome reveals extensive megaplasmid gene expression. **A)** A 109 kb region of the clade 1-like plasmid assembled and manually curated using transcripts encodes 11 protein coding genes with >500X transcript coverage. These include two pilin subunits (44, 45), VirB7 (51), IcmD/DotP (62), DotF (66), TPP1 (87), RuvB (89), and SlyB (94). Three highly expressed proteins have no functional predictions (33, 38, 90). The region also includes proteins of interest InvG (50) and FtsH (57). For the panda metagenome-derived reference, gene numbering is based on pMEGA4 homologs (genes 63 and 64, encoding transposon genes, are absent in the panda megaplasmid genome). **B**) Left - The predicted structure of two pilin subunit genes from pMEGA4 (44, 45), which share 92.4% amino acid identity. Right - The two structures aligned to *E. coli* major structural subunit of bundle-forming pilus BfpA (AF-P33553). **C)** An example region of transcript mapping to pMEGA3 showing protein coding genes 96, 97 and 105 with > 100X coverage. These are a protein related to telomerase-recruitment factor TPP1, a protein most similar to a Cas1-4 fusion, and SlyB. **D)** All of the 8 almost sequential proteins profiled as RNAse H1 in **B**have a domain that aligns well with PDB 1WSJ (red). **E)** Monomer of SlyB (pMEGA3_105; median pLDDT 90.06) and its alignment to PDB 7OJF.

Most genes are expressed at relatively even levels across both clade representatives. The most extreme peaks in RNA abundance are all associated with intergenic regions, two of which in the clade 2 genome are > 1 kb in length. Many of the highly expressed protein-coding genes lack functional assignments (49% and 55% of highly expressed proteins in clades 1 and 2, respectively), and some are very small (<100 amino acids; 23% in both; **Tables S20, S21**). However, proteins predicted to be involved with the T4SS and pilus are highly expressed in the clade 1-like genome, including two copies of a major pilin subunit (**Figure 6AB**). A region of pMEGA3 (**Figure 6C**) that features a large cluster of proteins with RNAse H1 (as well as other) domains (**Figure 6D**) includes a highly expressed protein annotated in part as TPP1, a possible Cas1-Cas4 fusion protein, and SlyB (**Figure 6E**). Also expressed but at lower levels are two Clp proteases and a L,D-transpeptidase YcbB, possibly involved in resistance to β-lactams due to alternative cross-linking of peptidoglycan^72^.

Members from 10 protein subfamilies are highly expressed by megaplasmids of both Clades, including SlyB, a protein involved in lipid biosynthesis, the ribosome hibernation factor YhbH, IcmD/DotP, tellurium resistance protein TerD, oxygen damage protein YfiD, TPP1, a Type IV pilin subunit, and two unknown proteins (**Tables S20, S21**). Of the 170 tRNAs in pMEGA3 and 108 in pMEGA4, virtually all recruited transcripts.

From the assembled transcripts, ribosomal protein S3 diversity revealed a highly abundant *E. coli* (∼1615X), another potential *E. coli* with lower coverage (∼33X), as well as multiple *Bacillota* (ie. *Clostridiales* and *Bacillales*) and a potential *Bacteroidota* member. Binning of transcript-assembled contigs to the *E. coli* OP50 genome (CP059137.1) resulted in 525 contigs representing 4.02 Mb, with 99.15% ANI over 79.04% of the OP50 genome. Given the presence of two megaplasmids and two members of *Enterobacteriaceae*, the high coverage Clade 1-like plasmid is likely hosted by the highly abundant *E. coli*, while the Clade 2-like plasmid may associate with the less abundant *E. coli* in the panda gut community.

## Discussion

We report the discovery and bioinformatic analysis of two clades of megaplasmids. The megaplasmids are clearly distinguished from bacteriophages by the absence of structural genes and from chromids/chromosomes by their low fraction of metabolic genes, protein novelty, plasmid-associated genes, replication style, and differences in GC content compared to host genomes. Members of both pMEGA clades whose genomes were reconstructed from isolates are confidently linked to *E. coli* and *Salmonella spp.* hosts. More broadly, analyses of protein similarity, co-occurrence, and DNA modification patterns indicate that the hosts are likely all *Enterobacteriaceae*. With genomes of up to 1.58 Mb, these are, by far, the largest megaplasmids in bacteria of this family.

While initially identified in preterm infant gut microbiomes, the megaplasmids occur in microbiomes of the human adult gut, animal gut, and other environments, suggesting they are widespread. All but one of the 20 genomes is “complete”, rather than bins or putatively circularized sequences. The exception is pMEGA3, which was reconstructed from PacBio reads but was too low in abundance to very 100% of the bases using Illumina reads. Complete genomes ensured accurate determination of genome lengths, provided information about genome organization and replication style, and facilitated accurate gene and proteome predictions. *In silico* structure prediction and analysis helped identify proteins implicated in virulence, components of the T4SS, proteins involved in host-cell adhesion, antibiotic resistance, and phage defense. Overall, the findings suggest that these megaplasmids are self-transmissible. Extensive methylation motifs to protect from diverse host restriction systems suggest that the pMEGAs may be able to replicate in multiple hosts.

Plasmids must ensure that the burden of their existence does not compromise the host organism’s viability^73^. The degree to which plasmids must offset the cost of their carriage is probably size-dependent and may drive plasmid acquisition of genes that increase the host’s fitness. Presumably, pMEGA genomes became large partly via the acquisition of gene variants from prior hosts. This likely occurred in an ancestral plasmid, as all pMEGAs have very low GC content compared to *Enterobacteriaceae* hosts (i.e., the GC content of host-derived genes has been ameliorated).

Presumably reflecting the acquisition of host functions, there is substantial overlap in gene content of pMEGA4 and its *E. coli* host. Some plasmid-borne variants may confer advantages by different specificity or enzyme kinetics, by supplying genes for stress protection, or by extending growth over a wider range of conditions. For example, plasmid-borne Clp and FtsH proteases exhibit different thermotolerance^74–77^ or specificities^48,49,78–80^ that may expand substrate range. Associated membrane microdomains (HflK/C) may result in compartmentalization of specific proteins involved in signal transduction, membrane trafficking, and regulation of proteases, including FtsH^51,52,81^. Other plasmid-derived benefits include expanding defense against invading bacteriophages and other plasmids (e.g., by provision of additional CRISPR-Cas systems). Plasmids may also increase antibiotic resistance by supplying efflux pumps with different specificities, remodelling the cell wall, and expressing variants of thymidylate synthase, LpxC, and ClpP that are insensitive to antibiotics^72,82–84^.

Megaplasmids may acquire functions to counter host defenses. They encode genes for NAD production, probably to offset decreased availability of NAD when the host depletes its NAD pool in response to infection^85^. Host tRNA degradation in response to invasion by ECEs may be countered by megaplasmid tRNA ligases, as has been shown for phages^86^. The pMEGAs encode upwards of 100 tRNAs. Although some certainly offset differences in codon use between the pMEGA and its host (**Figure 5**), structural diversity in tRNAs with the same anticodon may confer a spectrum of resistances to host tRNA nucleases. This counter-defense is well established in phages^87^. There is apparently strong selective pressure to preserve the codon and for sequence divergence outside of the codon.

Megaplasmids encode functions that do not duplicate those of their hosts. Of potential medical importance in the pMEGA4 plasmid in the ST10 *E. coli* strain is InvG. This protein forms the outer membrane (OM) basal body in the Type III secretion system injectisome that delivers effector proteins into host cells, but is also found in Type II secretion systems, Type IV pili structures, and phage assembly complexes^55,88,89^ (**Figure 4**). Putative toxin and poly-immunity arrays may enhance inter-bacterial competition and protect from diverse toxins, whereas diverse adhesion factors (ie. vWA domains) may contribute to motility and virulence^90,91^. Highly prevalent multicopy proteins have alpha- and beta-solenoid structures, which are often found in secreted effectors^92^. Effectors containing ankyrin repeats from intracellular pathogens such as *Legionella pneumophila* and *Coxiella burnetii* are involved in the invasion of eukaryotic cells^93^. Alpha-solenoid folds are more common in proteins of obligate intracellular pathogens as compared to other bacterial genomes^94^. Ankyrin repeat proteins (a form of alpha-solenoids) often mediate protein-protein interactions and may have functions analogous to those of DARPins, which are alpha-solenoids that are engineered to bind to target molecules with high affinity and specificity for functions such as antibodies and as anti-viral agents^95^. It is also possible that the megaplasmids accumulate ankyrin proteins to modulate their bacterial host proteome, or modulate eukaryotic immune response, as has been demonstrated in phage^96,97^.

Some megaplasmids associate with bacteria that cause gastroenteritis (i.e., *S. enterica*) and sepsis (i.e., *E. coli*)^98,99^. The pMEGA4 host *E. coli* is an ESBL-producing ST10 clone. ESBL-producing ST10 clones of *E. coli* have been increasing in prevalence globally^100^ and are common in animals, including chickens, associated food products, and humans^101–104^. Some other megaplasmid-bearing *E. coli* and *S.* Enteriditis isolates originate from human samples and one *E. coli* isolate of serotype O157:H7 was associated with an outbreak due to contaminated lettuce (Biosample: SAMN14113693)^105,106^. These observations motivate the investigation of the medical significance of megaplasmids.

## Conclusions

We identified the first megabase-scale plasmids in *Enterobacteriaceae*. They were detected in infants, adults and animals, as well as in hospital sinks where they likely serve as a reservoir for human transmission^107^. The megaplasmid genomes encode many solenoid proteins that may bind selectively to proteins or other molecules. We also identified mechanisms to extend host and habitat ranges, increase competitiveness in microbiomes, counter host defense, and protect from antibiotics. Much remains to be learned about their large and incompletely defined functional potential, but there are many indications that they hold medical significance. From the perspective of future work, it is important that they occur in genetically tractable *Enterobacteriaceae*, including *E. coli*.

## Acknowledgements

We thank Dr. Rani Phoolwanti for input regarding the RNAase H1 proteins and Colin Robinson for input regarding the metatranscriptome data. We thank the generators of the panda metatranscriptome data SRR10902853, whom, to date, we have not been able to identify. We thank Dr. Shufei Lei and Leylen Miloslavich for technical support. AKG, JFB, OTT, RS, and SW were supported by the Innovative Genomics Institute.

## Author Contributions

AKG and JFB were responsible for conceptualization and design of this study. Assembly was performed by AKG, and genome curation was performed by AKG and JFB. AKG, JFB, OTT, L-XC performed metagenome and metatranscriptome data analysis. SW and RS analysed the DNA modification patterns. OTT performed CRISPR analyses. NM and MH generated the nanopore sequencing of the *E. coli* isolate. AD, SB, VTD, AM, JC and AK provided data for megaplasmid detection. BF coordinated and submitted infant stool samples for sequencing. RS provided computational support. AJ, LS, and MM provided input regarding clinical relevance. Figures were generated by AKG, SW, OTT, and JFB. The manuscript was written by JFB and AKG, with input from OTT. JFB and MJM provided most of the funding support. All authors read and approved the manuscript prior to submission.

## Declaration of interests

The authors declare no conflicts of interest.

## Declaration of generative AI and AI-assisted technologies

No generative AI and AI-assisted technologies were used in the writing process.

## Funding

The primary funding for this project was provided by NIH award 5R01AI092531-14 to JFB, MM, with some support from the Chan Zuckerberg Foundation (CZIF2022-007203) via the Innovative Genomics Institute. Long-read PacBio sequencing was provided by a SMRT grant to AKG from PacBio and SeqCenter. RS and SW were funded by Lyda Hill Philanthropies, Acton Family Giving, the Valhalla Foundation, Hastings/Quillin Fund, the CH Foundation, Laura and Gary Lauder and Family, the Sea Grape Foundation, the Emerson Collective, Mike Schroepfer and Erin Hoffman Family Fund, the Anne Wojcicki Foundation. AHD and AEM were funded by the Biotechnology and Biological Sciences Research Council (BBSRC) Institute projects BB/X011011/1 and BBS/E/QU/230002A. AHD was funded by a UKRI BBSRC Doctoral Training Grant BB/T008717/1. AK was funded by the Finnish Multidisciplinary Centre of Excellence in Antimicrobial Resistance Research (FIMAR; 346127). MH received funding from the Swiss National Science Foundation grant IZJFZ3-177614.

## Data and code availability

Prior to publication, the genome sequences described in this study can be accessed and downloaded from ggKbase: https://ggkbase.berkeley.edu/project_groups/Megaplasmids_public. Raw sequencing data will be made available on NCBI upon publication. Other data reported in this study can be accessed at: 10.5281/zenodo.1723140.

## Methods

### DNA extraction, metagenomic sequencing, and binning

Stool samples from two preterm infants enrolled in the NICU Antibiotics and Outcomes (NANO) trial (Clinical Trial ID: NCT03997266) were collected over the first eight weeks of life after obtaining informed parental consent^108^. DNA was extracted from samples using the ZymoBIOMICS^TM^ DNA Miniprep Kit. Libraries prepared with the Illumina DNA prep kit were sequenced on an Illumina NovaSeq 6000 (2 x 151 bp reads). Reads were trimmed using sickle v1.33, and mapped against the human T2T reference using bowtie2 v2.5.4, resulting in an average of 4.09 gigabases per sample (**Table S1**)^109,110^. Metagenomic assembly was performed using metaSPAdes v4.0.0^111^. Scaffolds greater than 1000 bp were annotated using Prodigal v2.6.3, and searched against KEGG, UniProt, and UniRef using USearch v10.0.240 (-ublast)^112–116^. Ribosomal RNA sequences were predicted as previously described, and tRNA sequences were identified using tRNAscan-SE v1.3.1^117–119^.

For a subset of samples (**Table S1**), either the same DNA extract used for Illumina sequencing, or a new DNA extract prepared as above but with a modified lysis step was sequenced on a PacBio Revio. HiFi reads were trimmed using BBDuk v39.10 and assembled using hifiasm-meta v0.3-r073, resulting in, on average, 6.99 gigabases per sample^120,121^. Binning of Illumina short-read data to generate metagenomic-assembled genomes (MAGs) was accomplished using a combination of CONCOCT v1.1.0, MaxBin2 v2.2.7, MetaBAT2 v2.15 and Vamb v3.0.2, and the best bins were selected using DAS Tool v1.1.6^122^ with default settings^122–126^. The PacBio HiFi-MAG-Pipeline v3.3.2 was used with slight modifications to detect putative complete circular genomes (93% completeness) and bin other scaffolds into MAGs with less than 100 contigs^127^. Other settings included: a score threshold of 0.2 for DAS Tool, a minimum completeness of 70%, and a maximum contamination of 10%. Metagenomic assemblies were visualized and binned manually through an in-house platform ggKbase (https://ggkbase.berkeley.edu/). MAGs from both Illumina and PacBio assemblies for each infant were dereplicated using dRep v.3.4.5^128^ (with completeness (-comp) 75, contamination (-con) 10, secondary cluster ANI (-sa) 0.98, and coverage threshold (-nc) 0.25). This resulted in 27 MAGs for INF134001 (total 110) and 16 MAGs for 1330004 (total 72). The taxonomy of each MAG was determined using PhyloPhlAn v3.0.36^129^.

### Identification and manual curation of megaplasmids in the preterm infant gut

Examination of a PacBio assembly of one infant sample (INF1340011_TP8) identified a 1.58 Mb (pMEGA1) contig of unknown origin, with 3 bacterial single-copy genes, no ribosomal rRNA sequences, and many gene annotations characteristic of a plasmid. Two additional putatively complete plasmids were identified in other PacBio assemblies; a 1.33 Mb plasmid (pMEGA2) in INF1330004 at TP4 and a 1.37 Mb plasmid (pMEGA3) in INF1340011 at 4 weeks. Trimmed and filtered reads were mapped against the putative complete megaplasmids with BBMap v39.10^120^ (default settings with ambiguous=random). The alignment file was imported into Geneious Prime v2025.1.1, and reads were re-mapped to allow 0% mismatches to identify regions of possible variation or error. Reads were mapped again to allow 4% mismatches to the reference to correct possible single-nucleotide polymorphisms, insertions, deletions, or assembly errors. This process was repeated until no errors were visible in the plasmid genome. Reads that surpassed the ends of the sequence were used to extend the plasmid until a repeated sequence was present at the beginning and end of the plasmid, suggesting circularity. Paired reads that mapped at both ends of the sequence were also used as an indication of a circular plasmid, as described previously^130,131^. CoverM v0.7.0 was used along with Illumina reads to determine the depth and percentage of reads mapped to pMEGAs and the dereplicated bacterial MAGs for each infant across all available timepoints^132^.

### Identification and manual curation of a megaplasmid in *Escherichia coli*

To further assess the diversity and distribution of these megaplasmids in other datasets, we used the bacterial single-copy gene histidyl-tRNA synthetase (hisS) as a marker. A BlastP search against NCBI non-redundant protein sequences (executed December 16th, 2024) resulted in 1 highly similar hit for pMEGA1 (*Escherichia coli* – 93% cov, 97.63% aa ID, BioSample: SAMN18191027, Assembly: GCA_024673755.1)^133–135^. The top BlastP match for pMEGA2 is from a *Serratia sp.* with 98% coverage and 63.84% amino acid similarity (WP_282994782.1). The *E. coli* sequence is from a genomic assembly of an isolate (G269_1R) from Bangkok, Thailand^31,136^. We assembled the Illumina sequencing reads from this accession using the same methods as described above. Scaffolds that were distinct in GC content from the host genome (<30% for pMEGA vs ∼51% for *E. coli*), similar in read coverage, and that had no consensus for taxonomy based on the taxonomy of predicted gene sequences were binned. We also mapped scaffolds from the assembly to the complete pMEGA1 and pMEGA2 sequences from above using minimap2 v.2.28-r1209 (with the flag -x map-hifi)^137^. From these two approaches, we manually selected 23 scaffolds encompassing ∼1.39 Mb to represent a putative megaplasmid. Through this approach, and to the best of our ability and knowledge, we believe these scaffolds do not belong to *E. coli* or other easily identified mobile genetic elements such as other plasmids or phages.

Extensive manual curation and read mapping did not result in a complete circular element, so Nanopore long-read sequencing was performed on the *E. coli* isolate. Briefly, genomic DNA from the isolate was prepared with the Rapid sequencing DNA kit (SQK-RBK114.24) and sequenced on a GridION MKI R10 flow cell. Reads were basecalled with the high-accuracy model using Dorado v7.2.13^138^. Long-read sequencing resulted in very low coverage of the putative megaplasmid, but a complete 4.8 Mb genome for *E. coli* and a 104 kbp plasmid. The few Nanopore long-reads that were recovered against the 1.39 Mb plasmid were used to scaffold the contigs derived from Illumina sequencing. Through many iterative rounds of read mapping and extension from the ends of Illumina contigs and into divergent gaps where no direct read recruitment was possible, we reconstructed a complete 1.41 Mb plasmid (pMEGA4).

### Identification and manual curation of megaplasmids in other public datasets

We employed two tools to search for similar sequences across read data deposited in NCBI’s Sequence Read Archive (SRA) and other collections. First, PebbleScout was used to search against all available databases with the complete reference sequences of pMEGA1 and pMEGA2 elements^139^. This resulted in 55 hits with a PB score >60 for pMEGA1, and 319 hits with a PB score >60 for pMEGA2. Next, we searched all available assembled data in Logan (11-20-2024) using the *hisS* gene sequence (813 bp) from pMEGA1 and 1000 bp from the start of the *hisS* gene sequence from pMEGA2^140^. This resulted in 61 hits for pMEGA1 and almost 500 hits for pMEGA2 with a kmer coverage >=0.70. Many of the hits overlapped with the PebbleScout search results, therefore, we selected a few studies and any bacterial isolate hits only for additional investigation (**Table S5**). The bioprojects PRJEB36775 and PRJNA938144 relate to hospital sink metagenomes and infant gut metagenomes, respectively, and were selected given their high number of hits and relevance to human health^141,142^.

For all selected hits, reads from the SRA or European Nucleotide Archive (ENA) were downloaded and processed as described above for trimming and human read removal, if needed. Each read accession was treated as a separate sample regardless of whether more than one hit originated from the same BioSample (eg. hits from PRJEB21259 and PRJEB36775). Trimmed reads were mapped using bowtie2 and samtools to the pMEGA1 and pMEGA2 reference sequences to determine the percent length coverage, depth of coverage and percentage of reads mapped to the pMEGA in each sample^110,143^. We proceeded to assemble the hits using metaSPAdes v4.0.0 as described above and assessed to what extent complete sequences could be recovered by mapping scaffolds to pMEGA1 and pMEGA2 as references using minimap2^111,137^. Samples were considered a high-quality hit for a pMEGA if 10 or fewer scaffolds provided coverage of >=85% of the reference pMEGA. Samples were considered medium-quality hits if 11 to 50 scaffolds provided coverage of >=85% of a pMEGA, and all other hits were considered low-quality. From the high-quality hits, a selection of samples was chosen for manual curation as described above to attempt to complete the plasmid sequences and confirm circularity.

From the PebbleScout hits, SRR10902853 is listed as an RNA-seq dataset. To the best of our knowledge, this sample and its corresponding Bioproject (PRJNA601687) are not affiliated with a publication. We were unable to find an associated metagenomic dataset for this sample. We mapped the trimmed reads to a concatenated reference of pMEGA1,2,3 and 4 using bowtie2 v2.5.4 and visualized the alignments in Geneious Prime v2025.1.1 to assess regions with high coverage. For the clade 1-like plasmid, we assembled and manually curated a 109 kb fragment with >98% nucleotide identity to pMEGA4. A cut-off of 100X coverage was chosen to classify ORFs with high coverage for the clade 1-like genome, based on transcripts mapping to pMEGA3, and 500X coverage was chosen as a comparable level of high expression in the clade 2-like genome, based on transcripts mapping to pMEGA4 and the curated fragments. The clade 1-like plasmid had approximately 5X as many transcripts mapped as the clade 2-like plasmid. From the assembly, we detected ribosomal protein S3 sequences using hmmsearch with PF00189^144^. Contigs were mapped to the *E. coli* OP50 genome (CP059137.1) with minimap2, and ANI was calculated with skani^137,145^.

### Comparing the protein repertoire across 20 confident and circular megaplasmids

After curation, we recovered 20 circular megaplasmid sequences (**Table S7**). These sequences were re-oriented to start at a highly conserved protein sequence (ParE) and aligned using Mauve v1.1.3 in Geneious Prime^146^. Skani v0.2.1 and MASH v2.2 dist were used to assess the average nucleotide identity and distance between entire plasmid sequences^145,147^. The taxonomy of predicted proteins within these plasmid genomes was summarized using Uniprot annotations and tRep (https://github.com/MrOlm/tRep/tree/master/bin). All proteins across the 20 megaplasmids were clustered into protein subfamilies using MMseqs2 and an all-vs-all search (cluster workflow with E-value 0.001, sensitivity: 7.5, fraction of covered residues across target and query: 0.5, and the Set-Cover (greedy) cluster mode)^148^. A multiple sequence alignment was generated for subfamilies with at least 2 members using (mmseqs result2msa) and HMM profiles were generated using hhmake from HH-suite3^149^. A HH-Suite database was generated from these protein subfamilies and subfamilies were compared against each other using their respective HMMs and HHblits (with parameters -v 0 -p 50 -E 0.001 -z 1 -Z 32000 -B 0 -b 0 -n 2). Protein families were generated from subfamilies with probability scores of at least 95% and coverage of at least 0.70 using a Markov clustering algorithm^150^. Protein subfamilies were annotated based on HMM-HMM comparisons against PFAM using HHsearch v.3.0.3 (parameters -p 50 -E 0.001 -z 1 -Z 32000 -b 0 -B 0 -n 1)^151,152^.

### Additional annotation of megaplasmid features and protein structure prediction

Mobile genetic element prediction with geNomad v1.7.4 does not recognize the pMEGAs as plasmids but predicts proviruses within the genome^153^. More extensive annotation was performed for the pMEGA1-4 sequences including prediction of secretion systems and conjugation machinery using hmmscan against the CONJScan and TXSS hmm databases with an E value cut-off of 1e-10^154,155^. Origins of transfer (*oriT*) and vegetative replication (*oriV*) were predicted by oriTFinder2, oriTDB^34^ and OriV-Finder^156^. Repeats were identified manually in Geneious Prime using RepeatFinder. Virulence factors and toxins were predicted for pMEGA4 and the *E. coli* genome using PathoFact v.2.0^157^. We used PADLOC v2.0.0 and structural comparisons to identify antiviral defense systems and CRISPRCasFinder v.4.2.18 to detect CRISPR spacers^158,159^. Antibiotic resistance mechanisms in pMEGA4 were predicted from protein sequences using the *main* function of the Resistance Gene Identifier (RGI) v.6.0.3 with CARD v4.0.0 (settings: --include_loose, --include_nudge, --low_quality and reporting hits with a bitscore >100 regardless of ARO cutoff)^160^. For the *E. coli* genome and its 104 kb plasmid, RGI was used in the default mode with the --low_quality flag. Codon usage was calculated using coRdon v 1.26.0 in R v 4.4.3^161^.

For pMEGAs 1-4, AlphaFold2 was used to predict the protein structure of all open reading frames and compared to a database of folded proteins from PDB (retrieved 2023-04-20) using FoldSeek (v427df8a6b5d0ef78bee0f98cd3e6faaca18f172d)^162,163^. Some structures, as well as multimers, were predicted using AlphaFold3^162^. Multimer confidence was evaluated by consideration of ipTM scores. The top-scoring hits to PDB structures (generally >200 bit score) were manually visualized in ChimeraX (v1.9) and position-resolved pLDDT values were used to evaluate confidence. Alignment was performed using matchmaker or TM-align to confirm structural similarity^164,165^. Foldseek matches to structures in the AlphaFold database were also considered.

Multi-copy protein subfamilies were defined as subfamilies with greater than three members within at least one genome of pMEGA1-4. Multi-copy proteins and clusters of two or more proteins of the same subfamily were identified in Microsoft Excel and with custom scripts in R v 4.4.3. The predicted structures of members within a multi-copy protein subfamily were visualized to confirm their predicted function and to identify structures that contain alpha solenoid-like folds/domains. Alpha-solenoid domains were identified by visual inspection, in some cases flagged by best matches to DARPINs^42^. These proteins often had very low bit scores for best matches to proteins in PDB.

### Phylogeny of plasmid relaxase and replication initiation proteins

To generate a phylogeny of known MOB_F_ relaxase proteins, sequences were gathered from two comprehensive studies of conjugative plasmids and through an hmmscan for MOB_F_ sequences in PLSDB and IMG/PR, resulting in 1854 hits^33,155,166,167^. These were clustered at 99% identity using vsearch v2.13.3 (--cluster_fast) and combined with the predicted MOB_F_ relaxase sequences from pMEGA1-4 (SF1156 and SF0489), resulting in 881 sequences^168^. These sequences were aligned using MAFFT v7.505, then a phylogenetic tree was generated using FastTree v2.1.11, rooted at the mid-point, and visualized using iTOL. A representative structure of SF0489 (pMEGA4_1394 - median pLDDT 90.19) and SF1156 (pMEGA4_58 - median pLDDT 90.5) was aligned to RecD2 bound to ssDNA (PDB:3GP8)^169^. The TM alignment scores are 0.73 and 0.98, respectively. Similarly, a phylogeny of 105 replication initiation proteins from plasmids of known incompatibility groups (n=65)^170^ was generated with predicted Rep sequences from pMEGAs (SF1471 and SF0640) after clustering at 90% identity. Rep sequences from megaplasmids identified in this study were collapsed as clades. Select protein structures were compared to the pMEGA4 representative of SF640 and SF1471 using FoldSeek (bitscores as heatmaps on the right of the tree). If not available in the AlphaFold database, the structures of Rep proteins were predicted with AlphaFold3. For SF0640, pMEGA4_629 (median pLDDT 94.12) was aligned to RepC from the *Salmonella enteritidis* IncQ plasmid pBLST (AFDB: A0A1D8X722) with TM-align (score 0.86). For SF1471, pMEGA4_1423 (median pLDDT 91.5) was aligned to TrfA from the *Escherichia coli* IncP plasmid pRK2 (AFDB: P07676) with TM-align (score 0.57). Finally, Rep protein structures were compared to the Rep protein from E. coli F plasmid bound to an iteron sequence (PDB: 1REP). The TM-align score with 1REP for pMEGA4_629 was 0.68 and for pMEGA4_1423 was 0.70.

### Host prediction through abundance correlation, DNA modification analysis, and Hi-C linkage

From the searches of the SRA, we identified *E. coli* and *Salmonella spp.* isolates with evidence of a megaplasmid. For one *E. coli* isolate, it was re-grown, DNA was re-isolated, and sequenced via Nanopore confirming the presence of the 1.4 Mb plasmid (pMEGA4). For infants with longitudinal samples (pMEGA1, pMEGA2, pMEGA3), we measured the Pearson Correlation in R v. 4.4.3 between the percentage of reads mapped to the megaplasmid and all bacterial MAGs over multiple sampling timepoints. For isolates with evidence of a medium- or high-quality megaplasmid, plasmid and potential chromosomal contigs were separated and binned manually based on GC content, coverage and similarity to pMEGA1 or 2. The megaplasmid copy number (PCN) was estimated by comparing the mean depth of Illumina read coverage of the plasmid and draft isolate genome as determined with CoverM v0.7.0 using BBMap v39.10^120,132^ (**Table S5**).

To detect DNA methylation from PacBio data, we computed the mean inter-pulse duration (IPD) at each base across the mapped reads. We empirically estimated baseline IPD values for each k-mer by comparing to the mean IPD values of the kmer from the metagenome. Bases with significantly elevated IPD ratios are inferred to be modified. Subsequently, motif discovery is carried out using MotifMaker (SMRT Link v13.1.0.221970). We then scanned each contig to evaluate the modified fraction of detected motifs, ultimately generating a genome-wide methylation profile. In addition, to annotate restriction endonucleases (REases) and methyltransferases (MTases) in the assemblies, we employed the annotate_rm module of MicrobeMod with default parameters^171^. Finally, the ProxiMeta™ Metagenome Deconvolution Platform (Hi-C) from Phase Genomics (Seattle, USA) was used to detect Hi-C linkages on the week 4 stool samples from INF1330004 and INF1340011.

### Additional investigation into CRISPR spacers and possible hosts

CRISPR spacers detected with CRISPRCasFinder v.4.2.18 from all 20 confident megaplasmid genomes were searched against their respective metagenomes using blastn-short (-perc_identity 97, -qcov_hsp_perc 100)^133^. In addition, CRISPR spacers predicted within the 20 metagenomic samples from which the plasmid genomes originated were searched against the megaplasmid genomes to identify potential hosts. Finally, CRISPR spacers from the 20 megaplasmid genomes were compared to PLSDB and IMG/PR and IMG/VR representing over 772,529 plasmid and 15,722,824 viral sequences^21^. Spacer interaction networks were visualized in CytoScape v 3.10.3. Finally, to infer potential hosts based on spacer acquisition against pMEGAs, we blasted over 322 million predicted spacers from 580,000 bacterial genomes^38^ and ∼450,000 public metagenomes^39^ against pMEGA1-20 and evaluated spacers with 100% identity hits.

### Comparing to other plasmid databases

To ensure our megaplasmid elements had not been previously characterized, we searched for similar elements in, to the best of our knowledge, all relevant and known plasmid and viral sequence databases including PLSDB (v2024-05-31_v2 - 72,556 sequences), IMG/PR (699,973 sequences), IMG/VR (15,722,824 sequences), 226,194 plasmids predicted from the human gut ^130^, as well as datasets from two recent studies that characterized plasmids (n = 265,476) and viruses (n = 160,478) from infants^172,173^ using MASH dist with pMEGA1 and pMEGA2^147^. To ensure that no plasmids of similar size to the pMEGAs have been described in *Enterobacteriaceae*, we analyzed all plasmid sequences >1Mb from PLSDB and IMG/PR, and RefSeq *Enterobacteriaceae* genomes with secondary replicons >1Mb. Many of these plasmids were derived from isolates. If a plasmid was derived from a metagenomic sample, we annotated the taxonomy of the host based on ANI to other plasmids or consensus taxonomy of predicted proteins. Finally, to prevent inclusion of mis-annotated plasmids or secondary chromosomes, we used CheckM2 to remove sequences with 50% or greater bacterial genome completeness^174^. Predicted proteins were annotated following the same basic pipeline for pMEGAs, and the percent of hypothetical, unknown, or uncharacterized proteins as determined by UniRef annotations was counted for both the largest contig of each isolate assembly and the putative >1Mb plasmid.

## SUPPLEMENTARY FIGURES

**Figure S1:**
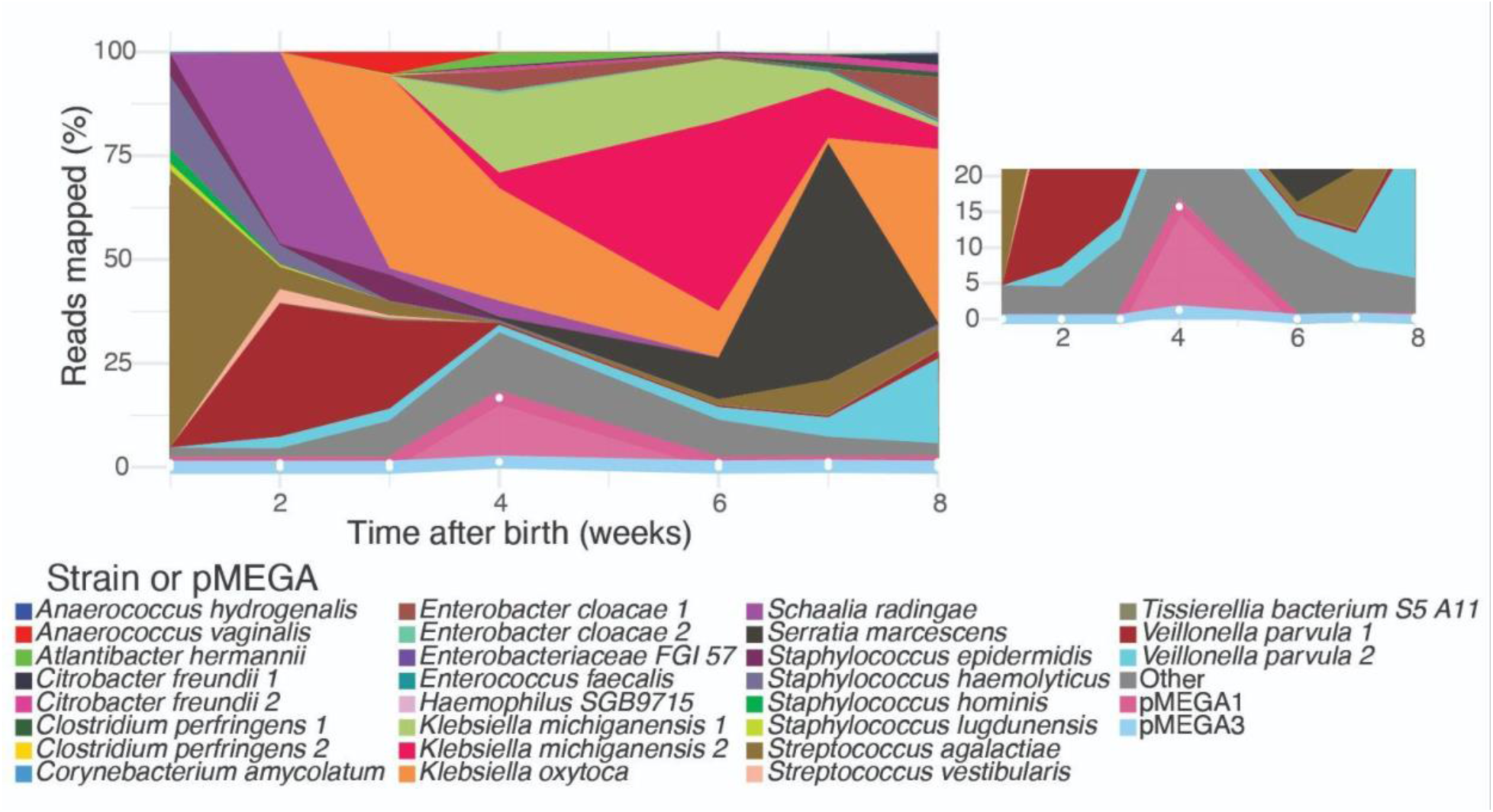
Infant gut microbiome community containing pMEGA1 and pMEGA3. The percentage of reads mapped to MAGs, pMEGA1 and pMEGA3 within INF1340011 across 8 weeks of life was determined through Illumina read mapping. The “Other” category includes reads that did not map to MAGs nor pMEGA1,3. Inset shows the same plot but with y values from 0 to 10.

**Figure S2:**
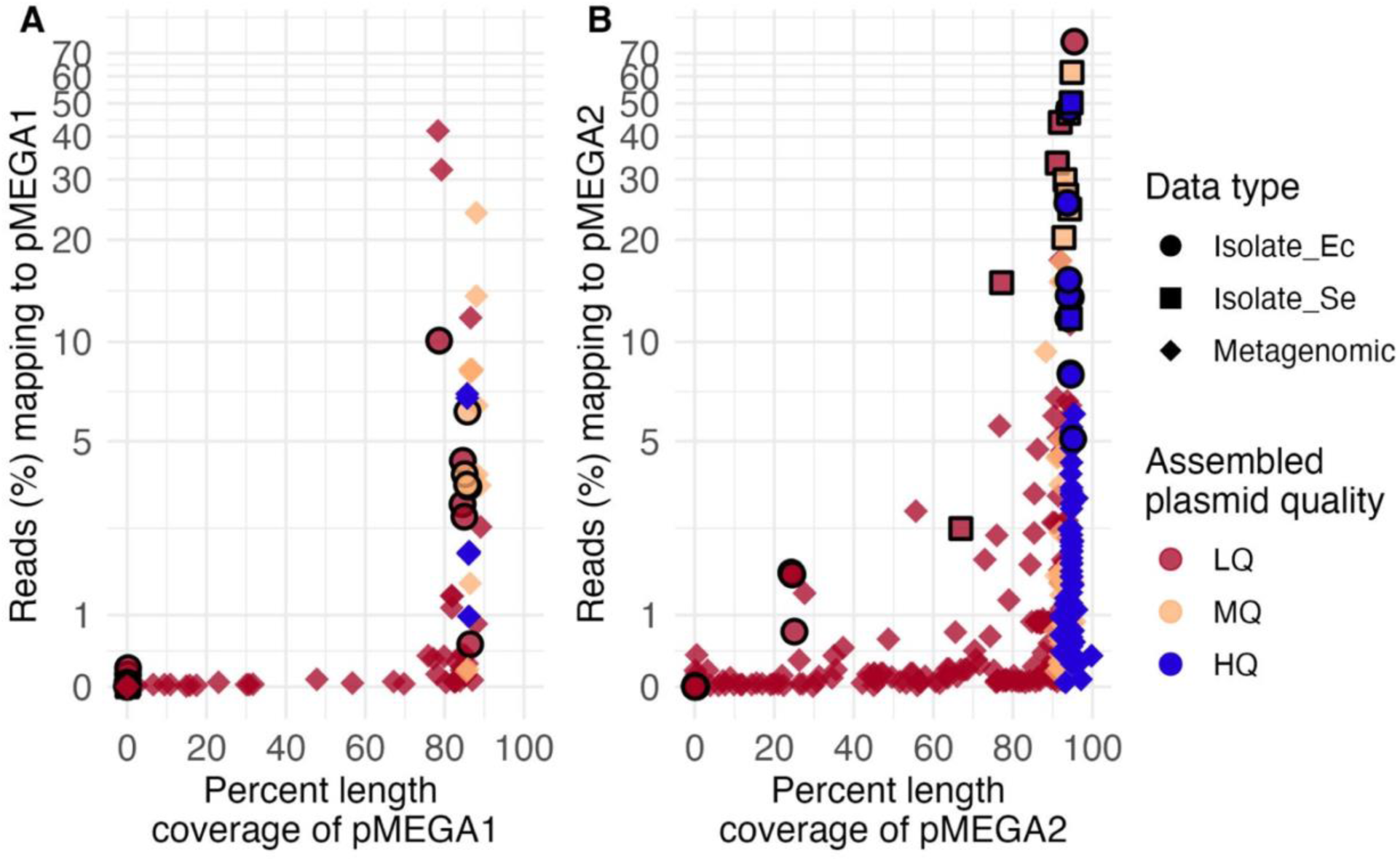
Prevalence and completeness of megaplasmids in public data. Sequencing reads from SRA accessions were mapped to pMEGA1 **(A)** and pMEGA2 **(B)** to determine the percent length coverage and percentage of reads mapped to each megaplasmid. For each accession, assembled plasmids were classified as high-quality (HQ - blue) if they had ≥85% coverage of a pMEGA and ≤10 contigs and medium-quality (MQ - yellow) with ≥85% coverage and 11-50 contigs. The remaining samples were classified as low-quality (LQ - red). Diamond: Metagenomic sample, circle: *Escherichia coli* Isolate (Ec), square: *Salmonella enteritidis* isolate (Se). Isolates are further highlighted by a black border.

**Figure S3:**
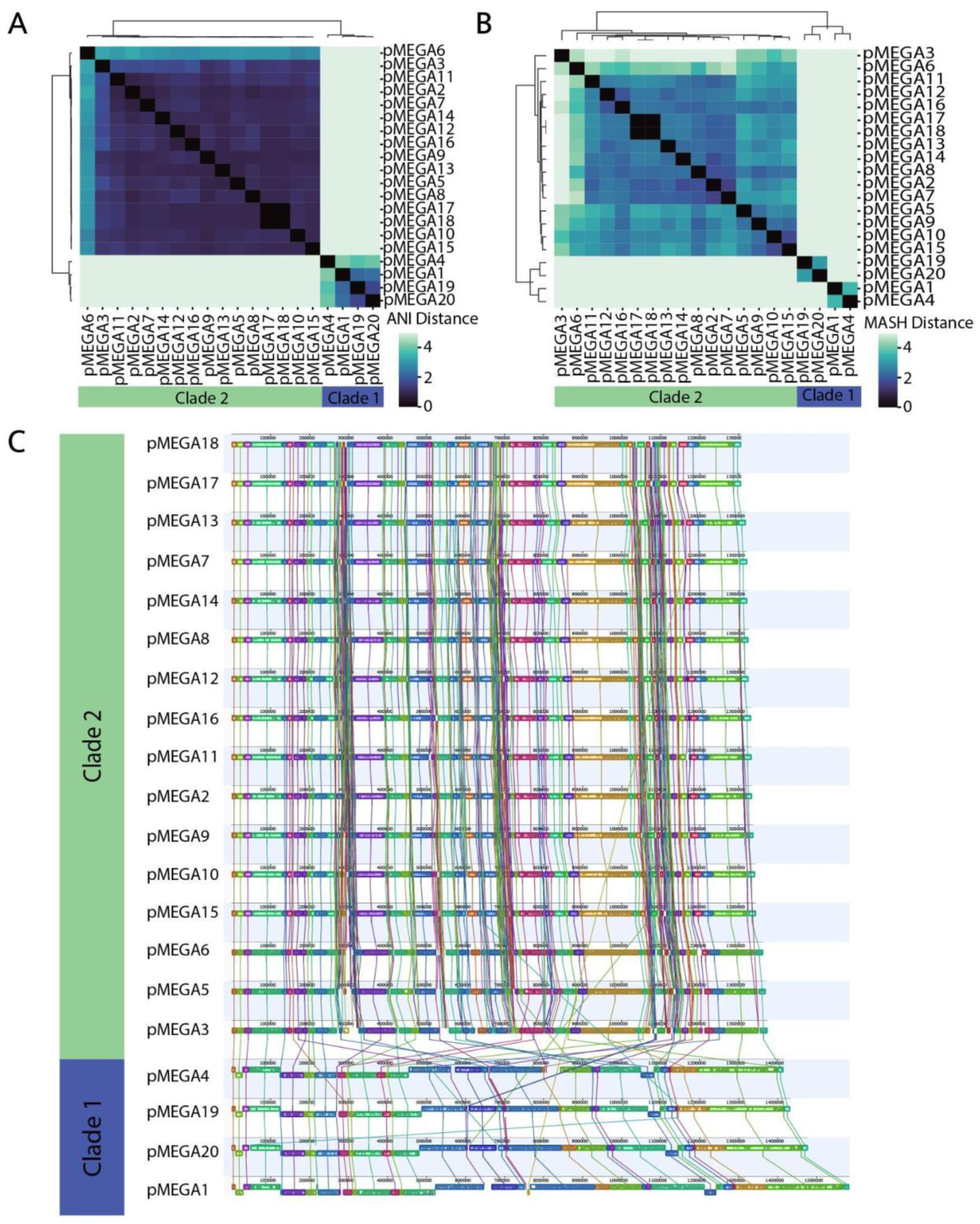
Comparing 20 complete megaplasmids. **A)** Whole genome alignment of complete plasmid sequences. Vertical lines connect blocks with similarity between plasmid genomes. The sequences were ordered manually and grouped based on clades. **B)** Matrix of average nucleotide identity (ANI). **C)** Matrix of MASH distance.

**Figure S4:**
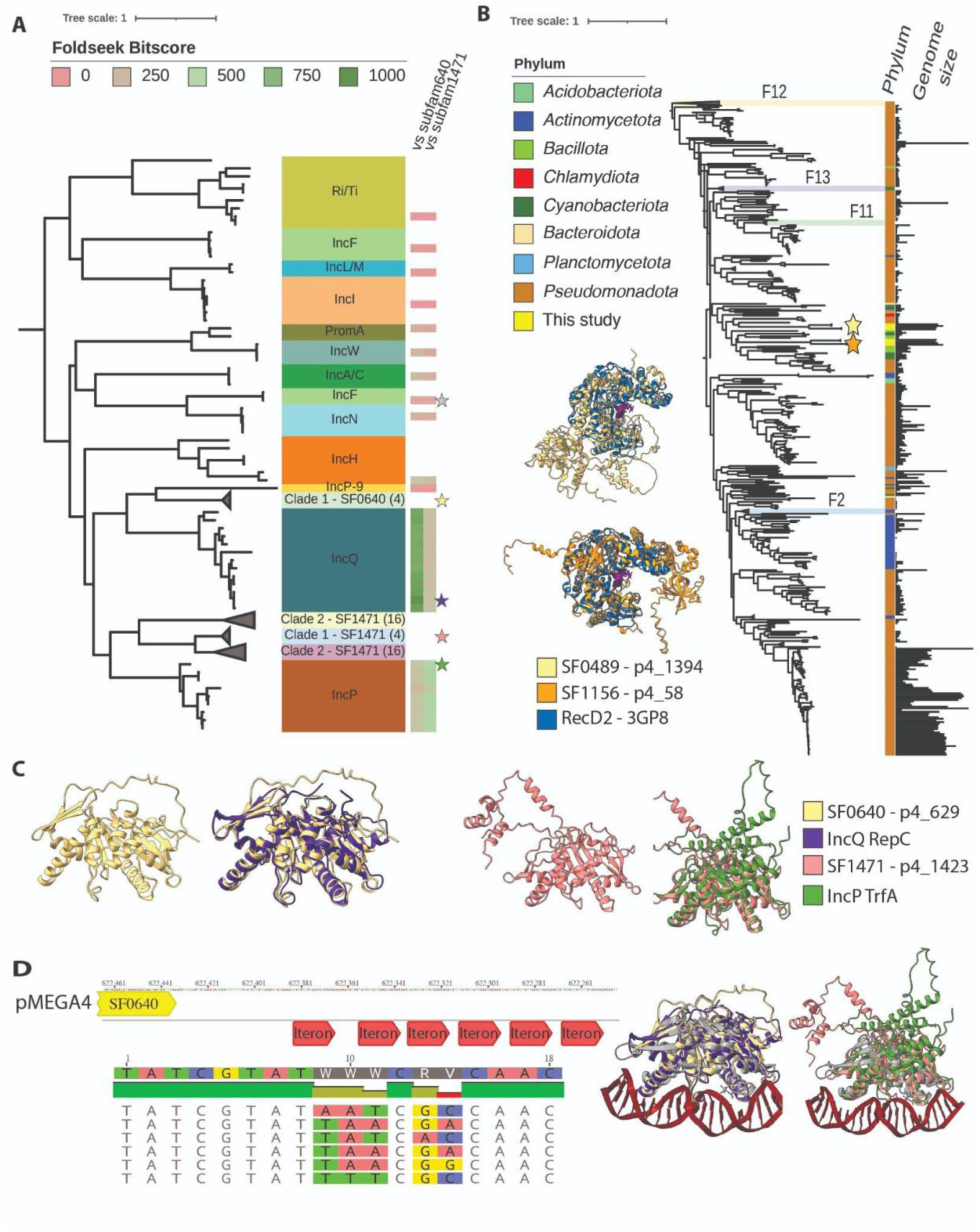
Replication initiation proteins and relaxases from two clades of megaplasmids. **A)** Phylogeny of 105 replication initiation (Rep) protein sequences. Reference Rep protein sequences are colored based on plasmid incompatibility group^170^. Bitscores from select FoldSeek hits are shown as heatmaps on the right of the tree. A star indicates that the protein structure is highlighted in panel C and D. **B)** Phylogeny of 881 MOB_F_ protein sequences with clades (F12, F13, F11, F2) collapsed^166^. Megaplasmids identified in this study are highlighted with a star. The length of each plasmid is shown on the far right panel. Predicted protein structures of relaxases aligned to PDB:3GP8 with ssDNA in purple are shown to the left of the tree. **C)** Representative structures of Rep proteins from pMEGA4 compared to RepC (purple) and TrfA (green). **D)** Region downstream of pMEGA4_629 with putative iterons and alignment of repeat sequences. To the right, alignment of structures from panel C to Rep protein from *E. coli* F plasmid (PDB: 1REP - grey) bound to the iteron sequence (red).

**Figure S5:**
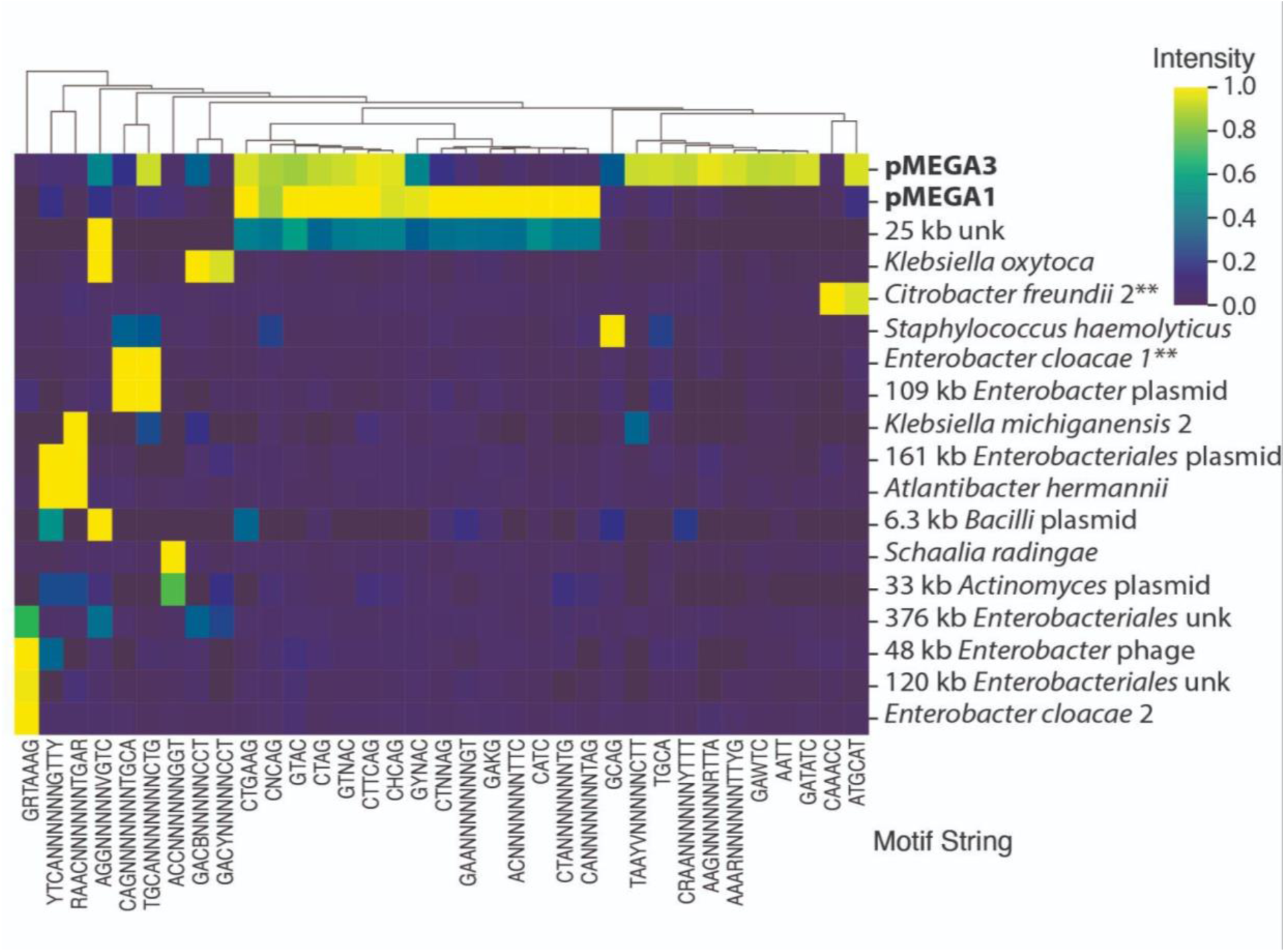
Possible *Enterobacteriaceae* hosts of pMEGA1 and pMEGA3. Modified bases were predicted in the PacBio HiFi reads of INF1340011 at week 4. The columns represent the motifs predicted to contain modified bases, and the rows represent genomes. ** Possible host(s) of pMEGA3 based on overlap in modified motif patterns.

**Figure S6:**
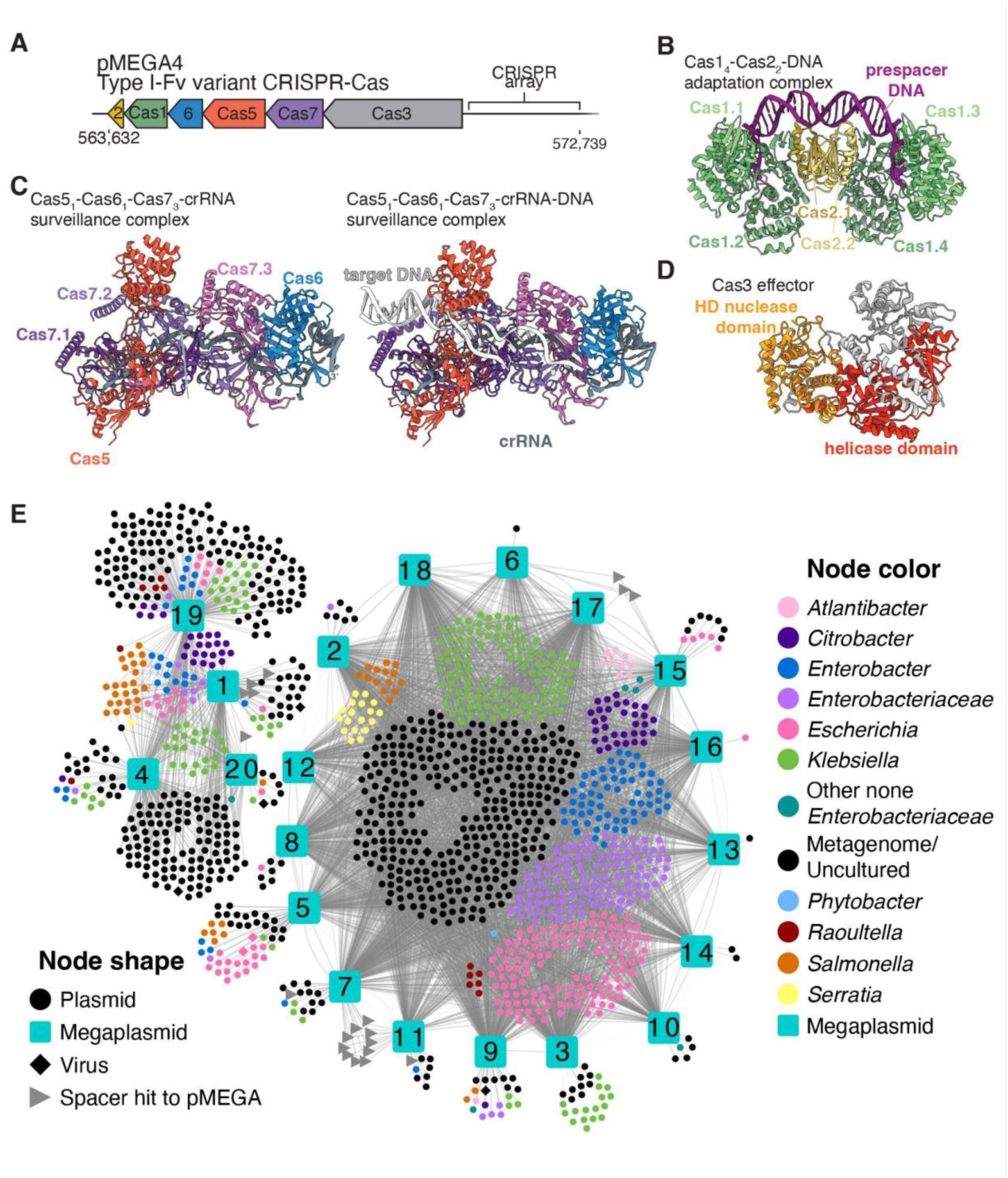
CRISPR-Cas system in pMEGA4 and targets of CRISPR spacers across pMEGA1-20. **A)** Diagram of the type I-Fv CRISPR-Cas locus from pMEGA4. **B)** Structural prediction of the pMEGA4 heterohexameric Cas1_4_-Cas2_2_ integrase complex (pMEGA4_568 and 569, median pLDDT 92.56 and 93.44) bound to a prespacer DNA. **C)** Structural predictions of the pMEGA4 Cascade surveillance complex in two states. Left, pMEGA4 Cas5_1_-Cas6_1_-Cas7_3_ (ORFs 570 - 572; median pLDDTs 85.19, 91.62, and 93.69) co-folded with a crRNA from an experimental structure (5O7H) of a similar type I-Fv system. Right, the same structure but co-folded with a target DNA. **D)** AlphaFold3 prediction of the pMEGA4 Cas3 effector (ORF 573 median pLDDT 91.81), with the HD nuclease domain colored in orange and the helicase domain colored in red. **E)** CRISPR spacer targets from pMEGA1-20 within other pMEGAs, plasmids from IMG/PR and PLSDB, and viral sequences from IMG/VR. Each line indicates a spacer targets a sequence in another element or, if the connection is between two pMEGA squares, the genomes share the same spacer with their CRISPR arrays. Grey triangles indicate a spacer from a pMEGA that targets another pMEGA.

**Figure S7:**
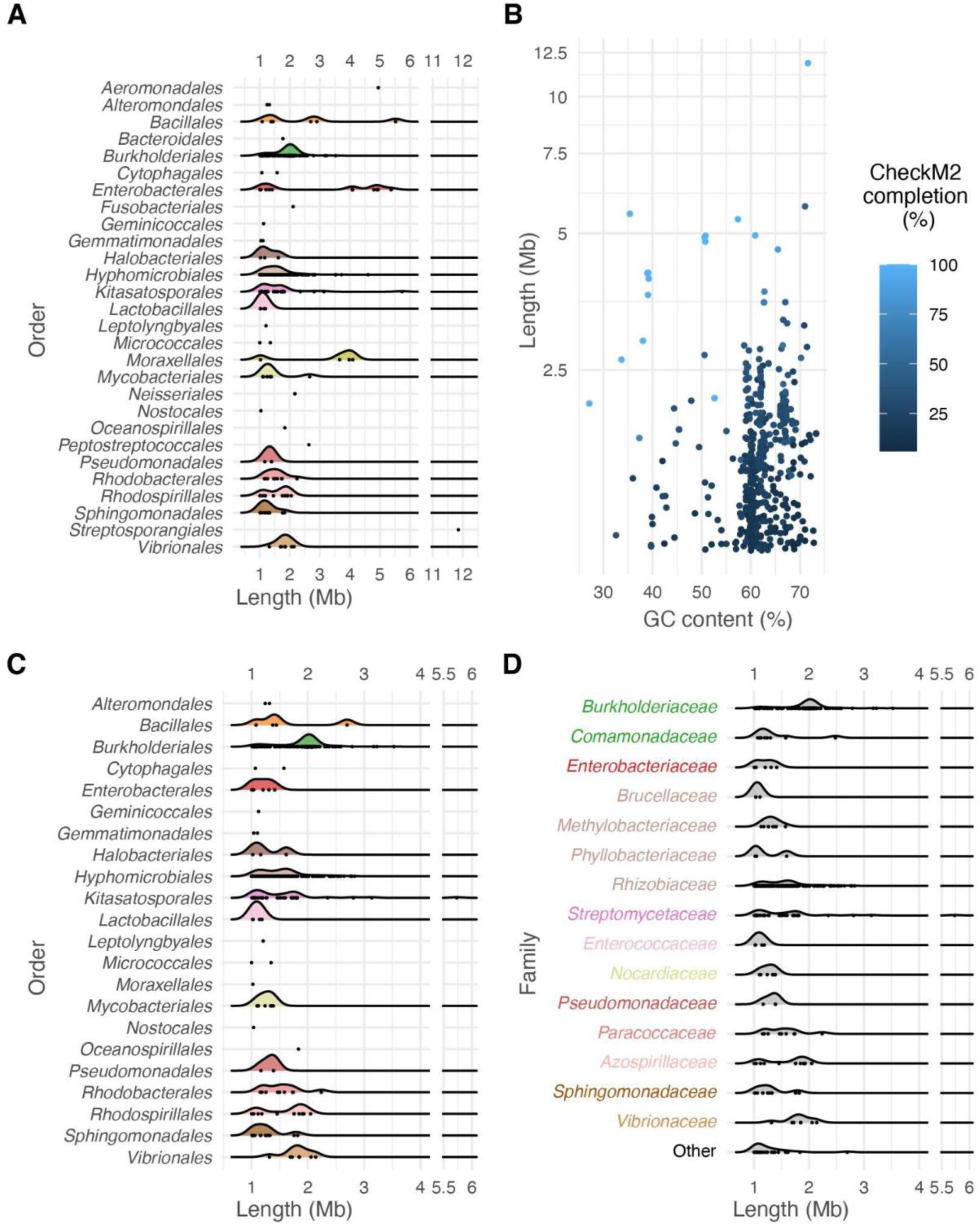
Plasmid databases are contaminated with complete bacterial genomes and possible secondary chromosomes. **A)** Length distribution of 667 non-redundant plasmids >1 Mb from PLSDB and IMG/PR grouped by the Order of their bacterial host. **B)** Length and GC content distribution of plasmid colored by completeness as determined with CheckM2. **C)** Length distribution of 642 plasmids >1Mb after removing sequences with >50% completion, grouped by taxonomic Order. **D)** Length distribution of 642 plasmids >1Mb after removing sequences with >50% completion, grouped by taxonomic Family. Families with two or fewer plasmids were grouped into Other.

**Figure S8:**
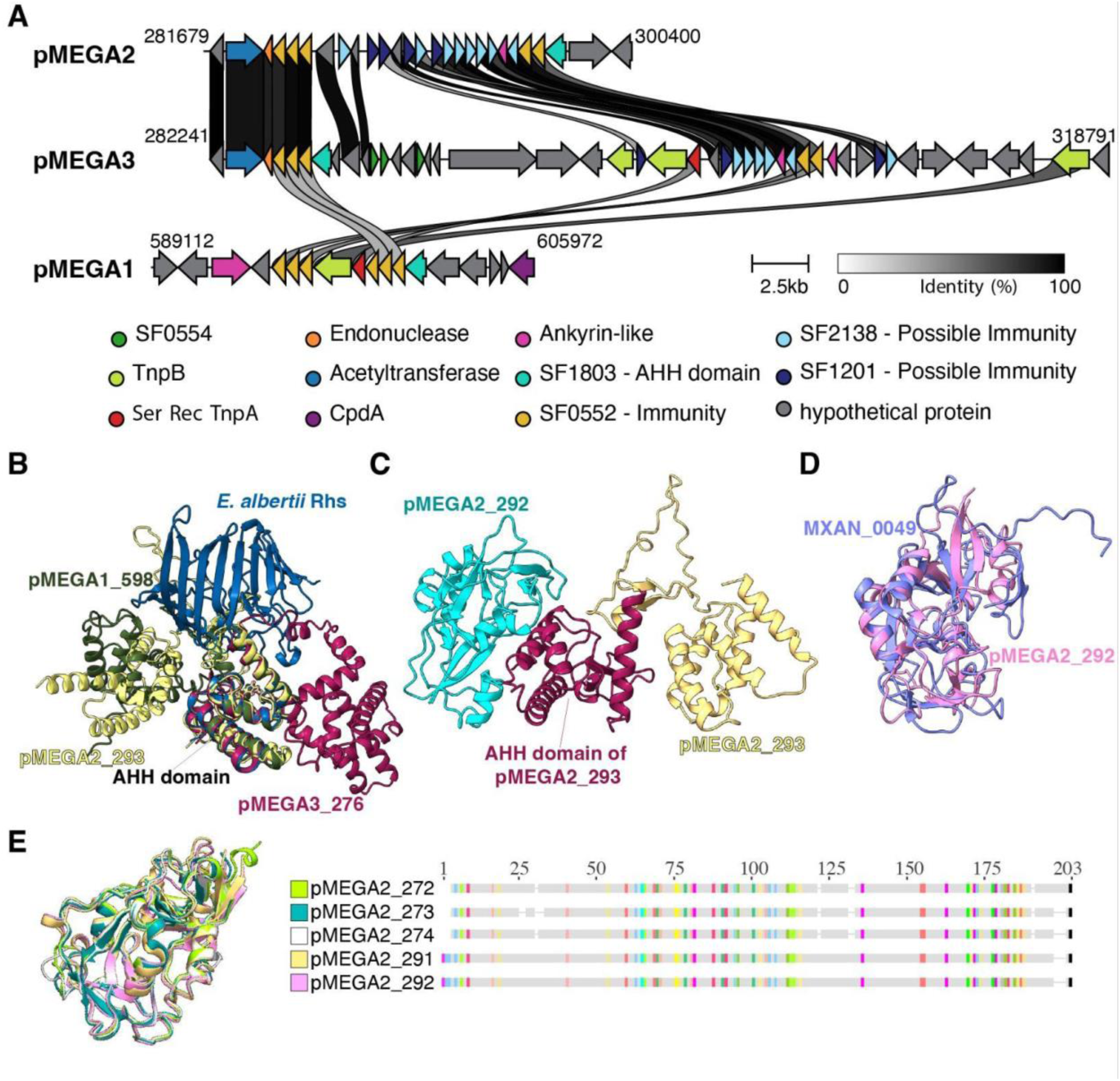
Polymorphic toxin and poly-immunity loci diversity in megaplasmids. **A)** Alignment of regions in pMEGAs1,2 and 3 with similar subfamilies, including a putative AHH nuclease toxin (SF1803) and tandem predicted immunity proteins (SF0552, 2138, 1201). **B)** Predicted polymorphic toxins of SF1803 (median pLDDTs: pMEGA1_598 71.88, pMEGA2_293 78.25, pMEGA3_276 69.12) aligned to a predicted structure of a Rhs family protein from *E. albertii* A0A855STS0 (blue). The aligned region is a predicted nuclease domain of the HNH/ENDO VII superfamily (PF14412) and contains a conserved AHH motif. **C)** Result of co-folding the predicted toxin AHH nuclease toxin with the downstream encoded predicted immunity protein. **D)** Alignment of the structure of a known nuclease immunity protein MXAN_0049 (PDB: 6A4C) to the predicted immunity protein (pink: pMEGA2_292 median pLDDT 96.38). **E)** Structural and protein sequence alignment of 5 members of SF0552 from pMEGA2.

**Figure S9:**
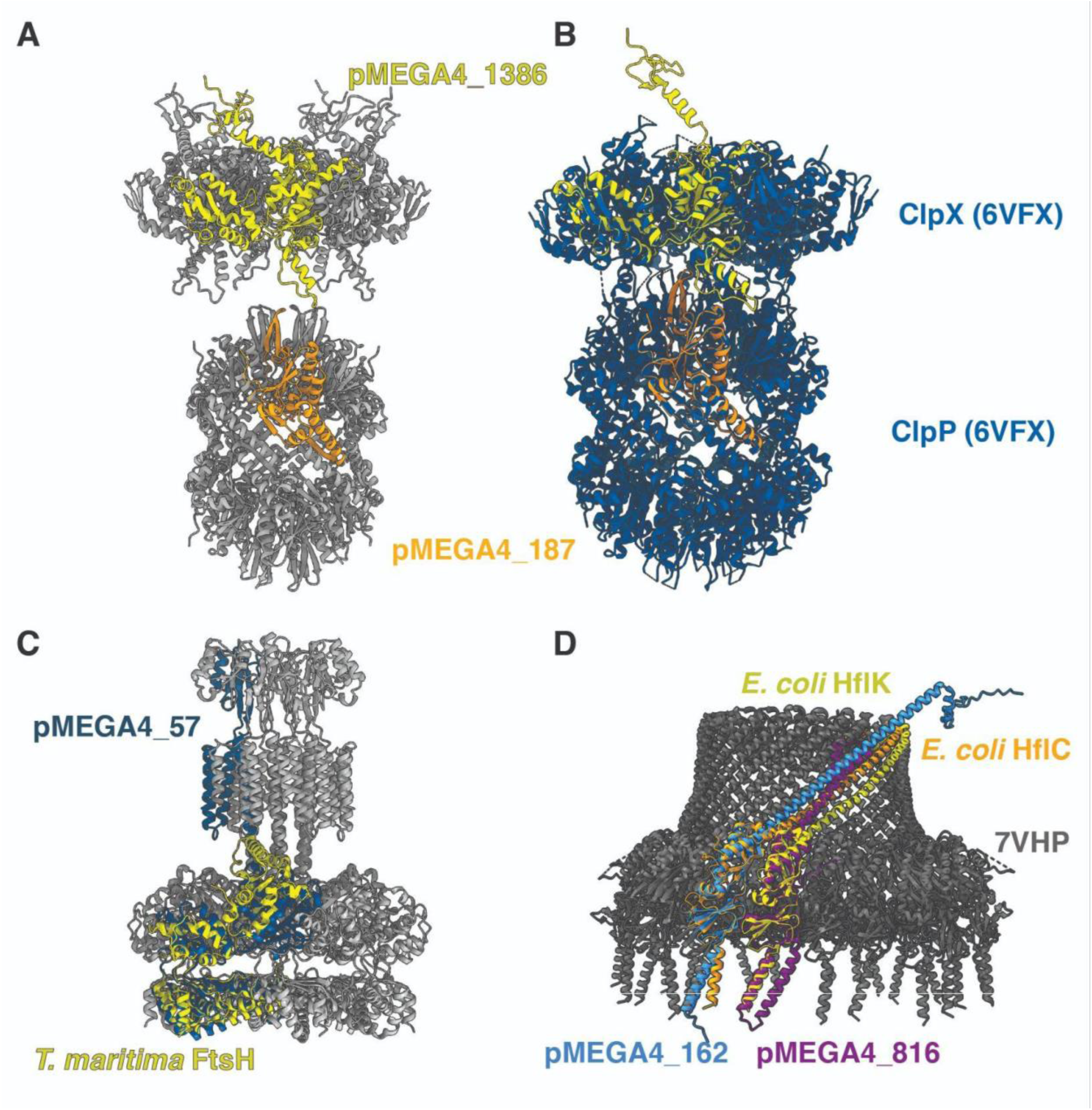
pMEGA4 predicted protein structures of various functions. **A)** Predicted hexamer of predicted ClpX (pMEGA4_1386) and dodecamer of ClpP (pMEGA4_187). **B)** Monomers of ClpX (median pLDDT 92.81) and ClpP (median pLDDT 98.56) from pMEGA4 aligned to reference of ClpXP from *Neisseria meningitidis* (6VFX). **C)** Predicted hexamer of FtsH (pMEGA4_57) in grey with monomer shown in blue and aligned to *Thermotoga maritima* FtsH (PDB: 3KDS). **D)** Predicted membrane microdomain proteins (median pLDDTs: pMEGA4_162 95.62, and pMEGA4_816 89.94) aligned to the structure of *E. coli* membrane microdomain (PDB: 7VHP). *E. coli* HflC and HflK are colored individually in orange and yellow.

## Supplementary Tables

**Table S1** List of NANO samples and assembly statistics

**Table S2** ANI and MASH distance between 20 complete plasmids

**Table S3** Relative abundance of MAGs and pMEGAs in INF1340011

**Table S4** Relative abundance of MAGs and pMEGAs in INF1330004

**Table S5** SRA read mapping and assembly from various studies

**Table S6** Hits from other internal NIH samples

**Table S7** List of 20 confident megaplasmids

**Table S8** pMEGA4 confident protein annotations

**Table S9** Core protein subfamilies identified across 20 megaplasmids

**Table S10** Correlations between MAG abundance and pMEGAs

**Table S11** Predicted modified motifs and methyltransferases within pMEGAs, other bacterial genomes, and MGEs in infant gut microbiome

**Table S12** Spacer hits to pMEGAs from isolate genomes and metagenomes

**Table S13** Putative Enterobacteriaceae 1Mb plasmids from NCBI

**Table S14** Multi-copy protein subfamilies and frequency of occurrence in clusters in pMEGA1-4

**Table S15** *E. coli* G269_1R ORF predictions

**Table S16** Comparing protein sequences from pMEGA4 to the *E. coli* host genome

**Table S17** tRNA sequence types in pMEGA4 and *E. coli* host

**Table S18** tRNA counts and codon usage in pMEGA4 and *E. coli* host

**Table S19** Antibiotic resistance genes predicted in *E. coli* G269_1R and pMEGA4

**Table S20** Highly expressed genes in clade 1-like megaplasmid from panda gut metatranscriptome dataset

**Table S21** Highly expressed genes in clade 2-like megaplasmid from panda gut metatranscriptome dataset

## Notes

https://doi.org/10.5281/zenodo.1723140

### Competing Interest Statement

The authors have declared no competing interest.

### Summary of Updates

Additional information on metatranscriptomic analysis added to methods and results; Figure 6 revised; author names corrected; grant number added; Supplementary tables 20 and 21 added

## References

1. Pilla, G. & Tang, C. M. Going around in circles: virulence plasmids in enteric pathogens. Nat. Rev. Microbiol. 16, 484–495 (2018).

2. Nora, L. C. et al. The art of vector engineering: towards the construction of next-generation genetic tools. Microb. Biotechnol. 12, 125–147 (2019).

3. Stockdale, S. R. & Hill, C. Incorporating plasmid biology and metagenomics into a holistic model of the human gut microbiome. Curr. Opin. Microbiol. 73, 102307 (2023).

4. Rodríguez-Beltrán, J., DelaFuente, J., León-Sampedro, R., MacLean, R. C. & San Millán, Á. Beyond horizontal gene transfer: the role of plasmids in bacterial evolution. Nat. Rev. Microbiol. 19, 347–359 (2021).

5. diCenzo, G. C. & Finan, T. M. The divided bacterial genome: Structure, function, and evolution. Microbiol. Mol. Biol. Rev. 81, (2017).

6. Maddamsetti, R. et al. Scaling laws of bacterial and archaeal plasmids. Nat. Commun. 16, (2025).

7. Ramiro-Martínez, P., de Quinto, I., Lanza, V. F., Gama, J. A. & Rodríguez-Beltrán, J. Universal rules govern plasmid copy number. Nat. Commun. 16, (2025).

8. Czarnecki, J., et al. Replication coordination marks the domestication of large extrachromosomal replicons in bacteria. bioRxiv (2025) doi:10.1101/2025.03.15.643453.

9. Genome sequence of the plant pathogen Ralstonia solanacearum. Nature (2002) doi:10.1038/news020128-8.

10. Janssen, P. J. et al. The complete genome sequence of Cupriavidus metallidurans strain CH34, a master survivalist in harsh and anthropogenic environments. PLoS One 5, e10433 (2010).

11. Mergeay, M. & Van Houdt, R. Cupriavidus metallidurans CH34, a historical perspective on its discovery, characterization and metal resistance. FEMS Microbiol. Ecol. 97, (2021).

12. diCenzo, G. C., Mengoni, A. & Perrin, E. Chromids aid genome expansion and functional diversification in the family Burkholderiaceae. Mol. Biol. Evol. 36, 562–574 (2019).

13. Fournes, F., Val, M.-E., Skovgaard, O. & Mazel, D. Replicate once per cell cycle: Replication control of secondary chromosomes. Front. Microbiol. 9, 1833 (2018).

14. Harrison, P. W., Lower, R. P. J., Kim, N. K. D. & Young, J. P. W. Introducing the bacterial “chromid”: not a chromosome, not a plasmid. Trends Microbiol. 18, 141–148 (2010).

15. Hall, J. P. J., Botelho, J., Cazares, A. & Baltrus, D. A. What makes a megaplasmid? Philos. Trans. R. Soc. Lond. B Biol. Sci. 377, 20200472 (2022).

16. Liu, H., et al. Unexplored diversity and potential functions of extra-chromosomal elements. mSystems e0017525 (2025).

17. Kohlmeier, M. G., O’Hara, G. W., Ramsay, J. P. & Terpolilli, J. J. Closed genomes of commercial inoculant rhizobia provide a blueprint for management of legume inoculation. Appl. Environ. Microbiol. 91, e0221324 (2025).

18. Medema, M. H. et al. The sequence of a 1.8-mb bacterial linear plasmid reveals a rich evolutionary reservoir of secondary metabolic pathways. Genome Biol. Evol. 2, 212–224 (2010).

19. Kothari, A. et al. Large circular plasmids from groundwater plasmidomes span multiple incompatibility groups and are enriched in multimetal resistance genes. MBio 10, (2019).

20. Redondo-Salvo, S. et al. Pathways for horizontal gene transfer in bacteria revealed by a global map of their plasmids. Nat. Commun. 11, 3602 (2020).

21. Schmartz, G. P. et al. PLSDB: advancing a comprehensive database of bacterial plasmids. Nucleic Acids Res. 50, D273–D278 (2022).

22. Camargo, A. P. et al. IMG/PR: a database of plasmids from genomes and metagenomes with rich annotations and metadata. Nucleic Acids Res. 52, D164–D173 (2024).

23. Li, Y. et al. PlasmidScope: a comprehensive plasmid database with rich annotations and online analytical tools. Nucleic Acids Res. 53, D179–D188 (2025).

24. Douarre, P.-E., Mallet, L., Radomski, N., Felten, A. & Mistou, M.-Y. Analysis of COMPASS, a new comprehensive Plasmid database revealed prevalence of multireplicon and extensive diversity of IncF plasmids. Front. Microbiol. 11, 483 (2020).

25. Koraimann, G. Spread and persistence of virulence and antibiotic resistance genes: A ride on the F Plasmid conjugation module. EcoSal Plus 8, (2018).

26. Ares-Arroyo, M., Rocha, E. P. C. & Gonzalez-Zorn, B. Evolution of ColE1-like plasmids across γ-Proteobacteria: From bacteriocin production to antimicrobial resistance. PLoS Genet. 17, (2021).

27. Arredondo-Alonso, S. et al. Plasmid-driven strategies for clone success in Escherichia coli. Nat. Commun. 16, 2921 (2025).

28. Brooks, B. et al. Strain-resolved analysis of hospital rooms and infants reveals overlap between the human and room microbiome. Nat. Commun. 8, 1814 (2017).

29. Shao, Y. et al. Stunted microbiota and opportunistic pathogen colonization in caesarean-section birth. Nature 574, 117–121 (2019).

30. Chen, Y. et al. Draft Genome Sequences of Citrobacter freundii and Citrobacter murliniae Strains Isolated from the Feces of Preterm Infants. Microbiol. Resour. Announc. 8, (2019).

31. Sudatip, D. et al. The risk of pig and chicken farming for carriage and transmission of Escherichia coli containing extended-spectrum beta-lactamase (ESBL) and mobile colistin resistance (mcr) genes in Thailand. Microb. Genom. 9, (2023).

32. Mishra, D. & Srinivasan, R. Catching a Walker in the act-DNA partitioning by ParA family of proteins. Front. Microbiol. 13, 856547 (2022).

33. Coluzzi, C., Garcillán-Barcia, M. P., de la Cruz, F. & Rocha, E. P. C. Evolution of Plasmid mobility: Origin and fate of conjugative and nonconjugative plasmids. Mol. Biol. Evol. 39, (2022).

34. Liu, G. et al. oriTDB: a database of the origin-of-transfer regions of bacterial mobile genetic elements. Nucleic Acids Res. 53, D163–D168 (2025).

35. Konieczny, I., Bury, K., Wawrzycka, A. & Wegrzyn, K. Iteron plasmids. Microbiol. Spectr. 2, (2014).

36. Smillie, C., Garcillán-Barcia, M. P., Francia, M. V., Rocha, E. P. C. & de la Cruz, F. Mobility of plasmids. Microbiol. Mol. Biol. Rev. 74, 434–452 (2010).

37. Pausch, P. et al. Structural variation of type I-F CRISPR RNA guided DNA surveillance. Mol. Cell 67, 622–632.e4 (2017).

38. Dion, M. B. et al. Streamlining CRISPR spacer-based bacterial host predictions to decipher the viral dark matter. Nucleic Acids Res. 49, 3127–3138 (2021).

39. Roux, S., et al. Planetary-scale metagenomic search reveals new patterns of CRISPR targeting. bioRxiv (2025) doi:10.1101/2025.06.12.659409.

40. Li, R., Peng, K., Li, Y., Liu, Y. & Wang, Z. Exploring tet(X)-bearing tigecycline-resistant bacteria of swine farming environments. Sci. Total Environ. 733, 139306 (2020).

41. Riccardi, C. et al. Independent origins and evolution of the secondary replicons of the class Gammaproteobacteria. Microb. Genom. 9, (2023).

42. Plückthun, A. Designed ankyrin repeat proteins (DARPins): binding proteins for research, diagnostics, and therapy. Annu. Rev. Pharmacol. Toxicol. 55, 489–511 (2015).

43. Gong, Y. et al. A nuclease-toxin and immunity system for kin discrimination in Myxococcus xanthus. Environ. Microbiol. 20, 2552–2567 (2018).

44. Zhang, D., Iyer, L. M. & Aravind, L. A novel immunity system for bacterial nucleic acid degrading toxins and its recruitment in various eukaryotic and DNA viral systems. Nucleic Acids Res. 39, 4532–4552 (2011).

45. Zhang, D., de Souza, R. F., Anantharaman, V., Iyer, L. M. & Aravind, L. Polymorphic toxin systems: Comprehensive characterization of trafficking modes, processing, mechanisms of action, immunity and ecology using comparative genomics. Biol. Direct 7, 18 (2012).

46. Ruhe, Z. C., Low, D. A. & Hayes, C. S. Polymorphic toxins and their immunity proteins: Diversity, evolution, and mechanisms of delivery. Annu. Rev. Microbiol. 74, 497–520 (2020).

47. Liu, W., Schoonen, M., Wang, T., McSweeney, S. & Liu, Q. Cryo-EM structure of transmembrane AAA+ protease FtsH in the ADP state. Commun. Biol. 5, 257 (2022).

48. Bittner, L.-M., Arends, J. & Narberhaus, F. When, how and why? Regulated proteolysis by the essential FtsH protease in Escherichia coli. Biol. Chem. 398, 625–635 (2017).

49. Erbse, A. et al. ClpS is an essential component of the N-end rule pathway in Escherichia coli. Nature 439, 753–756 (2006).

50. Deville, C., Franke, K., Mogk, A., Bukau, B. & Saibil, H. R. Two-step activation mechanism of the ClpB disaggregase for sequential substrate threading by the main ATPase motor. Cell Rep. 27, 3433–3446.e4 (2019).

51. Ma, C. et al. Structural insights into the membrane microdomain organization by SPFH family proteins. Cell Res. 32, 176–189 (2022).

52. Lopez, D. & Koch, G. Exploring functional membrane microdomains in bacteria: an overview. Curr. Opin. Microbiol. 36, 76–84 (2017).

53. Wallden, K., Rivera-Calzada, A. & Waksman, G. Type IV secretion systems: versatility and diversity in function. Cell. Microbiol. 12, 1203–1212 (2010).

54. Cascales, E. & Cambillau, C. Structural biology of type VI secretion systems. Philos. Trans. R. Soc. Lond. B Biol. Sci. 367, 1102–1111 (2012).

55. Hu, J. et al. Cryo-EM analysis of the T3S injectisome reveals the structure of the needle and open secretin. Nat. Commun. 9, 3840 (2018).

56. Hajam, I. A., Dar, P. A., Shahnawaz, I., Jaume, J. C. & Lee, J. H. Bacterial flagellin-a potent immunomodulatory agent. Exp. Mol. Med. 49, e373 (2017).

57. Haiko, J. & Westerlund-Wikström, B. The role of the bacterial flagellum in adhesion and virulence. Biology (Basel) 2, 1242–1267 (2013).

58. Ronneau, S. & Hallez, R. Make and break the alarmone: regulation of (p)ppGpp synthetase/hydrolase enzymes in bacteria. FEMS Microbiol. Rev. 43, 389–400 (2019).

59. Calhoun, L. N. & Kwon, Y. M. Structure, function and regulation of the DNA-binding protein Dps and its role in acid and oxidative stress resistance in Escherichia coli: a review. J. Appl. Microbiol. 110, 375–386 (2011).

60. Janssens, A. et al. SlyB encapsulates outer membrane proteins in stress-induced lipid nanodomains. Nature 626, 617–625 (2024).

61. Srikannathasan, V. et al. Structural basis for type VI secreted peptidoglycan DL-endopeptidase function, specificity and neutralization in Serratia marcescens. Acta Crystallogr. D Biol. Crystallogr. 69, 2468–2482 (2013).

62. Ting, S.-Y. et al. Bifunctional immunity proteins protect bacteria against FtsZ-targeting ADP-ribosylating toxins. Cell 175, 1380–1392.e14 (2018).

63. Bishop, R. E., Leskiw, B. K., Hodges, R. S., Kay, C. M. & Weiner, J. H. The entericidin locus of Escherichia coli and its implications for programmed bacterial cell death. J. Mol. Biol. 280, 583–596 (1998).

64. Schmidt, O. et al. prlF and yhaV encode a new toxin-antitoxin system in Escherichia coli. J. Mol. Biol. 372, 894–905 (2007).

65. Hansen, S. et al. Regulation of the Escherichia coli HipBA toxin-antitoxin system by proteolysis. PLoS One 7, e39185 (2012).

66. Tripathi, A., Dewan, P. C., Ahmed, S. & Varadarajan, R. MazF-induced growth inhibition and persister generation in Escherichia coli. J. Biol. Chem. 289, 4191–4205 (2014).

67. May, J. P. & Simon, A. E. Targeting of viral RNAs by Upf1-mediated RNA decay pathways. Curr. Opin. Virol. 47, 1–8 (2021).

68. Millman, A., et al. An expanding arsenal of immune systems that protect bacteria from phages. bioRxiv (2022) doi:10.1101/2022.05.11.491447.

69. Vassallo, C. N., Doering, C. R., Littlehale, M. L., Teodoro, G. I. C. & Laub, M. T. A functional selection reveals previously undetected anti-phage defence systems in the E. coli pangenome. Nat. Microbiol. 7, 1568–1579 (2022).

70. Gao, L. et al. Diverse enzymatic activities mediate antiviral immunity in prokaryotes. Science 369, 1077–1084 (2020).

71. Fitzpatrick, A. W. P. et al. Structure of the MacAB–TolC ABC-type tripartite multidrug efflux pump. Nat. Microbiol. 2, 17070 (2017).

72. Caveney, N. A. et al. Structural insight into YcbB-mediated beta-lactam resistance in Escherichia coli. Nat. Commun. 10, 1849 (2019).

73. San Millan, A. & MacLean, R. C. Fitness costs of plasmids: A limit to Plasmid transmission. Microbiol. Spectr. 5, (2017).

74. Kamal, S. M. et al. A recently isolated human commensal Escherichia coli ST10 clone member mediates enhanced thermotolerance and tetrathionate respiration on a P1 phage-derived IncY plasmid. Mol. Microbiol. 115, 255–271 (2021).

75. Lee, C. et al. Why? - Successful Pseudomonas aeruginosa clones with a focus on clone C. FEMS Microbiol. Rev. 44, 740–762 (2020).

76. Bojer, M. S., Struve, C., Ingmer, H., Hansen, D. S. & Krogfelt, K. A. Heat resistance mediated by a new plasmid encoded Clp ATPase, ClpK, as a possible novel mechanism for nosocomial persistence of Klebsiella pneumoniae. PLoS One 5, e15467 (2010).

77. Motiwala, T. et al. Caseinolytic proteins (Clp) in the genus Klebsiella: Special focus on ClpK. Molecules 27, 200 (2021).

78. Kamal, S. M. et al. Two FtsH proteases contribute to fitness and adaptation of Pseudomonas aeruginosa clone C strains. Front. Microbiol. 10, 1372 (2019).

79. Mawla, G. D. et al. The membrane-cytoplasmic linker defines activity of FtsH proteases in Pseudomonas aeruginosa clone C. J. Biol. Chem. 300, 105622 (2024).

80. Stein, B. J., Grant, R. A., Sauer, R. T. & Baker, T. A. Structural basis of an N-degron adaptor with more stringent specificity. Structure 24, 232–242 (2016).

81. Qiao, Z. et al. Cryo-EM structure of the entire FtsH-HflKC AAA protease complex. Cell Rep. 39, 110890 (2022).

82. Choi, M., Karunaratne, K. & Kohen, A. Flavin-dependent thymidylate synthase as a new antibiotic target. Molecules 21, 654 (2016).

83. Bacterial Enzymes as Targets for Drug Discovery. (Academic Press, San Diego, CA, 2024).

84. Zhao, J. et al. Preclinical safety and efficacy characterization of an LpxC inhibitor against Gram-negative pathogens. Sci. Transl. Med. 15, eadf5668 (2023).

85. Osterman, I. et al. Phages reconstitute NAD+ to counter bacterial immunity. Nature 634, 1160–1167 (2024).

86. Kaufmann, G. et al. RloC: A translation-disabling tRNase implicated in phage exclusion during recovery from DNA damage. in DNA Repair (InTech, 2011).

87. van den Berg, D. F., van der Steen, B. A., Costa, A. R. & Brouns, S. J. J. Phage tRNAs evade tRNA-targeting host defenses through anticodon loop mutations. Elife 12, (2023).

88. Korotkov, K. V., Gonen, T. & Hol, W. G. J. Secretins: dynamic channels for protein transport across membranes. Trends Biochem. Sci. 36, 433–443 (2011).

89. Leighton, T. L., Buensuceso, R. N. C., Howell, P. L. & Burrows, L. L. Biogenesis of Pseudomonas aeruginosa type IV pili and regulation of their function. Environ. Microbiol. 17, 4148–4163 (2015).

90. Hoffmann, L. et al. Structure and interactions of the archaeal motility repression module ArnA-ArnB that modulates archaellum gene expression in Sulfolobus acidocaldarius. J. Biol. Chem. 294, 7460–7471 (2019).

91. Heidler, T. V., Ernits, K., Ziolkowska, A., Claesson, R. & Persson, K. Porphyromonas gingivalis fimbrial protein Mfa5 contains a von Willebrand factor domain and an intramolecular isopeptide. Commun. Biol. 4, 106 (2021).

92. Patel, D. T. et al. Global atlas of predicted functional domains in Legionella pneumophila Dot/Icm translocated effectors. Mol. Syst. Biol. 21, 59–89 (2025).

93. Al-Khodor, S., Price, C. T., Habyarimana, F., Kalia, A. & Abu Kwaik, Y. A Dot/Icm-translocated ankyrin protein of Legionella pneumophila is required for intracellular proliferation within human macrophages and protozoa. Mol. Microbiol. 70, 908–923 (2008).

94. Schmitz-Esser, S. et al. The genome of the amoeba symbiont “Candidatus Amoebophilus asiaticus” reveals common mechanisms for host cell interaction among amoeba-associated bacteria. J. Bacteriol. 192, 1045–1057 (2010).

95. Walser, M., Mayor, J. & Rothenberger, S. Designed ankyrin repeat proteins: A new class of viral entry inhibitors. Viruses 14, 2242 (2022).

96. Al-Khodor, S., Price, C. T., Kalia, A. & Abu Kwaik, Y. Functional diversity of ankyrin repeats in microbial proteins. Trends Microbiol. 18, 132–139 (2010).

97. Jahn, M. T. et al. A phage protein aids bacterial symbionts in eukaryote immune evasion. Cell Host Microbe 26, 542–550.e5 (2019).

98. Self, J. L. et al. Epidemiology of salmonellosis among infants in the United States: 1968-2015. Pediatrics 151, (2023).

99. Hoffman, A. et al. Predictors of mortality and severe illness from Escherichia coli sepsis in neonates. J. Perinatol. 44, 1816–1821 (2024).

100. Yang, J.-T., Zhang, L.-J., Lu, Y., Zhang, R.-M. & Jiang, H.-X. Genomic insights into global blaCTX-M-55-positive Escherichia coli epidemiology and transmission characteristics. Microbiol. Spectr. 11, e0108923 (2023).

101. Day, M. J. et al. Extended-spectrum β-lactamase-producing Escherichia coli in human-derived and foodchain-derived samples from England, Wales, and Scotland: an epidemiological surveillance and typing study. Lancet Infect. Dis. 19, 1325–1335 (2019).

102. Falgenhauer, L. et al. Detection and characterization of ESBL-producing Escherichia coli from humans and poultry in Ghana. Front. Microbiol. 9, 3358 (2018).

103. Lupo, A., Saras, E., Madec, J.-Y. & Haenni, M. Emergence of blaCTX-M-55 associated with fosA, rmtB and mcr gene variants in Escherichia coli from various animal species in France. J. Antimicrob. Chemother. 73, 867–872 (2018).

104. Hu, Y.-Y. et al. Molecular typing of CTX-M-producing escherichia coli isolates from environmental water, swine feces, specimens from healthy humans, and human patients. Appl. Environ. Microbiol. 79, 5988–5996 (2013).

105. Lim, J. Y., Yoon, J. & Hovde, C. J. A brief overview of Escherichia coli O157:H7 and its plasmid O157. J. Microbiol. Biotechnol. 20, 5–14 (2010).

106. Chen, J. C. et al. Reoccurring Escherichia coli O157:H7 strain linked to leafy greens-associated outbreaks, 2016-2019. Emerg. Infect. Dis. 29, 1895–1899 (2023).

107. Kizny Gordon, A. E., et al. The hospital water environment as a reservoir for carbapenem-resistant organisms causing hospital-acquired infections-A systematic review of the literature. Clin. Infect. Dis. 64, 1435–1444 (2017).

108. Morowitz, M. J. et al. The NICU Antibiotics and Outcomes (NANO) trial: a randomized multicenter clinical trial assessing empiric antibiotics and clinical outcomes in newborn preterm infants. Trials 23, 428 (2022).

109. Joshi NA, F. J. N. Sickle: A sliding-window, adaptive, quality-based trimming tool for FastQ files (Version 1.33) [Software]. https://github.com/najoshi/sickle (2011).

110. Langmead, B. & Salzberg, S. L. Fast gapped-read alignment with Bowtie 2. Nat. Methods 9, 357–359 (2012).

111. Nurk, S., Meleshko, D., Korobeynikov, A. & Pevzner, P. A. metaSPAdes: a new versatile metagenomic assembler. Genome Res. 27, 824–834 (2017).

112. Hyatt, D. et al. Prodigal: prokaryotic gene recognition and translation initiation site identification. BMC Bioinformatics 11, 119 (2010).

113. Kanehisa, M. & Goto, S. KEGG: kyoto encyclopedia of genes and genomes. Nucleic Acids Res. 28, 27–30 (2000).

114. Suzek, B. E., Huang, H., McGarvey, P., Mazumder, R. & Wu, C. H. UniRef: comprehensive and non-redundant UniProt reference clusters. Bioinformatics 23, 1282–1288 (2007).

115. UniProt Consortium. UniProt: The universal protein knowledgebase in 2023. Nucleic Acids Res. 51, D523–D531 (2023).

116. Edgar, R. C. Search and clustering orders of magnitude faster than BLAST. Bioinformatics 26, 2460–2461 (2010).

117. Brown, C. T. et al. Unusual biology across a group comprising more than 15% of domain Bacteria. Nature 523, 208–211 (2015).

118. Nawrocki, E. P., Kolbe, D. L. & Eddy, S. R. Infernal 1.0: inference of RNA alignments. Bioinformatics 25, 1713–1713 (2009).

119. Lowe, T. M. & Eddy, S. R. tRNAscan-SE: a program for improved detection of transfer RNA genes in genomic sequence. Nucleic Acids Res. 25, 955–964 (1997).

120. Bushnell, B. BBMap.

121. Feng, X., Cheng, H., Portik, D. & Li, H. Metagenome assembly of high-fidelity long reads with hifiasm-meta. Nat. Methods 19, 671–674 (2022).

122. Sieber, C. M. K. et al. Recovery of genomes from metagenomes via a dereplication, aggregation and scoring strategy. Nat. Microbiol. 3, 836–843 (2018).

123. Wu, Y.-W., Simmons, B. A. & Singer, S. W. MaxBin 2.0: an automated binning algorithm to recover genomes from multiple metagenomic datasets. Bioinformatics 32, 605–607 (2016).

124. Alneberg, J. et al. Binning metagenomic contigs by coverage and composition. Nat. Methods 11, 1144–1146 (2014).

125. Nissen, J. N. et al. Improved metagenome binning and assembly using deep variational autoencoders. Nat. Biotechnol. 39, 555–560 (2021).

126. Kang, D. D. et al. MetaBAT 2: an adaptive binning algorithm for robust and efficient genome reconstruction from metagenome assemblies. PeerJ 7, e7359 (2019).

127. Portik, D. M., et al. Highly accurate metagenome-assembled genomes from human gut microbiota using long-read assembly, binning, and consolidation methods. bioRxiv (2024) doi:10.1101/2024.05.10.593587.

128. Olm, M. R., Brown, C. T., Brooks, B. & Banfield, J. F. dRep: a tool for fast and accurate genomic comparisons that enables improved genome recovery from metagenomes through de-replication. ISME J. 11, 2864–2868 (2017).

129. Asnicar, F. et al. Precise phylogenetic analysis of microbial isolates and genomes from metagenomes using PhyloPhlAn 3.0. Nat. Commun. 11, 2500 (2020).

130. Yu, M. K., Fogarty, E. C. & Eren, A. M. Diverse plasmid systems and their ecology across human gut metagenomes revealed by PlasX and MobMess. Nat. Microbiol. 9, 830–847 (2024).

131. Chen, L.-X., Anantharaman, K., Shaiber, A., Eren, A. M. & Banfield, J. F. Accurate and complete genomes from metagenomes. Genome Res. 30, 315–333 (2020).

132. Aroney, S. T. N. et al. CoverM: read alignment statistics for metagenomics. Bioinformatics 41, (2025).

133. Altschul, S. F., Gish, W., Miller, W., Myers, E. W. & Lipman, D. J. Basic local alignment search tool. J. Mol. Biol. 215, 403–410 (1990).

134. Bethesda (MD): National Library of Medicine (US), National Center for Biotechnology Information. Protein [Internet]. (1988).

135. Sayers, E. W. et al. Database resources of the National Center for Biotechnology Information in 2025. Nucleic Acids Res. 53, D20–D29 (2025).

136. Sudatip, D. et al. A One Health approach to assessing occupational exposure to antimicrobial resistance in Thailand: The FarmResist project. PLoS One 16, e0245250 (2021).

137. Li, H. Minimap2: pairwise alignment for nucleotide sequences. Bioinformatics 34, 3094–3100 (2018).

138. Nanoporetech. Dorado. (2024).

139. Shiryev, S. A. & Agarwala, R. Indexing and searching petabase-scale nucleotide resources. Nat. Methods 21, 994–1002 (2024).

140. Chikhi, R., Raffestin, B., Korobeynikov, A., Edgar, R. & Babaian, A. Logan: Planetary-scale genome assembly surveys life’s diversity. bioRxiv (2024) doi:10.1101/2024.07.30.605881.

141. Constantinides, B. et al. Genomic surveillance of Escherichia coli and Klebsiella spp. in hospital sink drains and patients. Microb. Genom. 6, (2020).

142. Zhang, A., et al. Infants exposed in utero to Hurricane Maria have gut microbiomes with reduced diversity and altered metabolic capacity. mSphere 8, e0013423 (2023).

143. Li, H. et al. The Sequence Alignment/Map format and SAMtools. Bioinformatics 25, 2078–2079 (2009).

144. Potter, S. C. et al. HMMER web server: 2018 update. Nucleic Acids Res. 46, W200–W204 (2018).

145. Shaw, J. & Yu, Y. W. Fast and robust metagenomic sequence comparison through sparse chaining with skani. Nat. Methods 20, 1661–1665 (2023).

146. Darling, A. C. E., Mau, B., Blattner, F. R. & Perna, N. T. Mauve: multiple alignment of conserved genomic sequence with rearrangements. Genome Res. 14, 1394–1403 (2004).

147. Ondov, B. D. et al. Mash: fast genome and metagenome distance estimation using MinHash. Genome Biol. 17, (2016).

148. Steinegger, M. & Söding, J. MMseqs2 enables sensitive protein sequence searching for the analysis of massive data sets. Nat Biotechnol 35, 1026–1028 (2017).

149. Steinegger, M. et al. HH-suite3 for fast remote homology detection and deep protein annotation. BMC Bioinformatics 20, 473 (2019).

150. Enright, A. J., Van Dongen, S. & Ouzounis, C. A. An efficient algorithm for large-scale detection of protein families. Nucleic Acids Res. 30, 1575–1584 (2002).

151. Söding, J. Protein homology detection by HMM-HMM comparison. Bioinformatics 21, 951–960 (2005).

152. Mistry, J. et al. Pfam: The protein families database in 2021. Nucleic Acids Res. 49, D412–D419 (2021).

153. Camargo, A. P. et al. Identification of mobile genetic elements with geNomad. Nat. Biotechnol. 42, 1303–1312 (2024).

154. Abby, S. S. et al. Identification of protein secretion systems in bacterial genomes. Sci. Rep. 6, 23080 (2016).

155. Cury, J., Abby, S. S., Doppelt-Azeroual, O., Néron, B. & Rocha, E. P. C. Identifying conjugative plasmids and integrative conjugative elements with CONJscan. Methods Mol. Biol. 2075, 265–283 (2020).

156. Li, Y. & Gao, F. OriV-Finder: a comprehensive web server for bacterial plasmid replication origin analysis. Nucleic Acids Res. 53, W451–W456 (2025).

157. Sunyer, J. O. et al. PathoFact 2.0: An integrative pipeline for predicting antimicrobial resistance genes, virulence factors, toxins and biosynthetic gene clusters in metagenomes. bioRxiv (2024) doi:10.1101/2024.12.09.627531.

158. Couvin, D. et al. CRISPRCasFinder, an update of CRISRFinder, includes a portable version, enhanced performance and integrates search for Cas proteins. Nucleic Acids Res. 46, W246–W251 (2018).

159. Payne, L. J. et al. Identification and classification of antiviral defence systems in bacteria and archaea with PADLOC reveals new system types. Nucleic Acids Res. 49, 10868–10878 (2021).

160. Alcock, B. P. et al. CARD 2023: expanded curation, support for machine learning, and resistome prediction at the Comprehensive Antibiotic Resistance Database. Nucleic Acids Res. 51, D690–D699 (2023).

161. Vlahovicek, A. E. M. K. CoRdon. (Bioconductor, 2018). doi:10.18129/B9.BIOC.CORDON.

162. Abramson, J. et al. Accurate structure prediction of biomolecular interactions with AlphaFold 3. Nature 630, 493–500 (2024).

163. van Kempen, M. et al. Fast and accurate protein structure search with Foldseek. Nat. Biotechnol. 42, 243–246 (2024).

164. Meng, E. C. et al. UCSF ChimeraX: Tools for structure building and analysis. Protein Sci. 32, e4792 (2023).

165. Zhang, Y. & Skolnick, J. TM-align: a protein structure alignment algorithm based on the TM-score. Nucleic Acids Res. 33, 2302–2309 (2005).

166. Garcillán-Barcia, M. P., Francia, M. V. & de la Cruz, F. The diversity of conjugative relaxases and its application in plasmid classification. FEMS Microbiol. Rev. 33, 657–687 (2009).

167. Eddy, S. R. Accelerated profile HMM searches. PLoS Comput. Biol. 7, e1002195 (2011).

168. Rognes, T., Flouri, T., Nichols, B., Quince, C. & Mahé, F. VSEARCH: a versatile open source tool for metagenomics. PeerJ (2016) doi:10.7287/peerj.preprints.2409.

169. Saikrishnan, K., Powell, B., Cook, N. J., Webb, M. R. & Wigley, D. B. Mechanistic basis of 5’-3’ translocation in SF1B helicases. Cell 137, 849–859 (2009).

170. Suzuki, H., Yano, H., Brown, C. J. & Top, E. M. Predicting plasmid promiscuity based on genomic signature. J. Bacteriol. 192, 6045–6055 (2010).

171. Crits-Christoph, A., Kang, S. C., Lee, H. H. & Ostrov, N. MicrobeMod: A computational toolkit for identifying prokaryotic methylation and restriction-modification with nanopore sequencing. bioRxiv (2023) doi:10.1101/2023.11.13.566931.

172. He, W., Russel, J., Klincke, F., Nesme, J. & Sørensen, S. J. Insights into the ecology of the infant gut plasmidome. Nat. Commun. 15, 6924 (2024).

173. Zeng, S. et al. A metagenomic catalog of the early-life human gut virome. Nat. Commun. 15, 1864 (2024).

174. Chklovski, A., Parks, D. H., Woodcroft, B. J. & Tyson, G. W. CheckM2: a rapid, scalable and accurate tool for assessing microbial genome quality using machine learning. Nat. Methods 20, 1203–1212 (2023).

175. Kreutzberger, M. A. B. et al. Convergent evolution in the supercoiling of prokaryotic flagellar filaments. Cell 185, 3487–3500.e14 (2022).

