## Supplementary Information for "Megaplasmids associate with *Escherichia coli* and other *Enterobacteriaceae*"

**Supplementary files and data are available at:** 10.5281/zenodo.1723140

**Other putative megaplasmids in *Enterobacteriaceae* should be classified as secondary chromosomes**

To expand on the information provided in Table S13, we describe some notable features of each secondary replicon >1 Mb in *Enterobacteriaceae* from NCBI.

Accessions with secondary replicons labelled as plasmids:

1. *Escherichia coli* strain STEC1012 plasmid pSTEC1012_1, accession number **CP061270**, has three rRNA genes, genes for citrate metabolism, cytochrome oxidase subunits, for enterobactin biosynthesis, folate metabolism, fimbrial production, heme biosynthesis, amino acid biosynthesis, lipopolysaccharide biosynthesis, molybdopterin biosynthesis, and some T4SS proteins. 19.87% of the proteins are hypothetical or contain only domains of unknown function.
2. *Escherichia coli* strain STEC411 plasmid pSTEC411_1, accession **CP061244.1** encodes two rRNAs, genes for cytochrome maturation, flagellar biosynthesis, many genes indicative of a prophage, metabolism (ethanolamine). Few plasmid annotations: plasmid partitioning/stability family protein CDS, plasmid segregation protein ParM CDS. 27.28% of the proteins have hypothetical or unknown function.
3. *Escherichia coli* strain STEC639 plasmid pSTEC639_1, accession **CP061233.1**, encodes 12 rRNAs (including the 16S and 23S rRNA genes), genes for amino acid

biosynthesis, metabolic genes, transporters, and genes for nitrate reduction. 23.44% of the proteins are hypothetical or contain domains of unknown function.

4. *Klebsiella pneumoniae* subsp. pneumoniae strain WRC19_AI1572C plasmid pAI1527P_P1, accession number **CP079635** does not have rRNAs but has numerous metabolic genes for processes including nitrate reduction, cytochrome biosynthesis, for type IV secretion, nucleotide biosynthesis, assimilation of hypoxanthine, cytochrome oxidase, many transporters, and likely a prophage. 33.20% of proteins are hypothetical or contain only domains of unknown function.
5. *Klebsiella pneumoniae* isolate 392 genome is a contig-level assembly. The putative plasmid P1, accession **OW848780.1,** does not have rRNAs but has ribosomal proteins, genes for cytochrome biosynthesis, for nitrate reduction, and many transporters. 28.59% of proteins hypothetical or contain domains of unknown function.
6. The linear element in *Klebsiella pneumoniae* strain LA86 plasmid pLAhvKp86-2, accession **CP182886**, is likely a genome fragment and is not terminated in inverted repeats. There are no annotated rRNAs. The fragment encodes numerous transporters, genes for nitrogen metabolism, many genes for phenolacetate degradation, catabolism of aromatic compounds, PQQ biosynthesis, nucleotide biosynthesis, and uric acid degradation. 24.22% of the genes are hypothetical or contain only domains of unknown function.
7. *Klebsiella pneumoniae* strain LAhvKp142 plasmid pLAhvKp142-1, accession **CP183011**, does not have rRNAs but encodes ribosomal proteins. The sequence encodes transporters, proteins involved in fatty acid metabolism, other carbohydrate metabolism, amino acid biosynthesis, and secretion systems. 23.20% of the proteins are hypothetical or contain only domains of unknown function.

The other accessions are unnamed or labelled numerically.

8. *Escherichia coli* strain RHB23-SO-C02 unnamed1 accession **CP099174** encodes three rRNAs (including the 16S and 23S rRNA genes), numerous genes for lipopolysaccharide, amino acid, and flagellar biosynthesis, and contains a well defined

prophage. 19.04% of proteins are hypothetical or contain only domains of unknown function.

9. *Salmonella enterica* strain SA88 unnamed1 accession **CP169339** encodes genes for lipopolysaccharide, amino acid, and flagellar biosynthesis, hydrogenases, and contains a prophage. It does not contain rRNAs. Of the 1702 genes annotated, 29.12% are hypothetical or contain only domains of unknown function.
10. *Salmonella enterica* subsp. enterica serovar Enteritidis strain 32 plasmid unnamed1, accession **CP183533** does not have annotations available from NCBI. We predicted this sequence contains 16S rRNAs, ribosomal proteins, and 30.04% of the proteins are hypothetical or contain domains of unknown function.
11. *Klebsiella aerogenes* strain NCTC10006 chromosome 6, accession LR134126, is part of an assembly that appears very fragmented. The largest contig (chromosome 3) is 1.49 Mb and is listed as circular. It is likely this genome is incomplete and elements are falsely circularized. Chromosome 6 contains rRNAs and ribosomal proteins. Only 14.92% of proteins are hypothetical or contain domains of unknown function.
12. For the remaining accessions (OZ185710.1,OZ185743.1,OZ185757.1) both the longest and secondary elements are labelled as linear. 26.8%, 22.71% and 26.73% of proteins have hypothetical or unknown annotations.
